# The Lassa Virus Fusion Domain has structural plasticity and exploits bis(monoacylglycero)phosphate for fusion

**DOI:** 10.1101/2025.05.20.655079

**Authors:** Hallie N. Pennington, Kiruthika Arulmoly, Quinn M. Mulvihill, Sungjai Shin, Wonpil Im, Jinwoo Lee

## Abstract

Infection with Lassa virus (LASV), an arenavirus endemic to West Africa, results in a viral hemorrhagic fever with high mortality rates and public health implications. The glycoprotein complex (GPC) is central to LASV’s infectivity as it mediates viral entry via membrane fusion. The fusion domain (FD, G^260^ – N^295^) facilitates the initiation of membrane fusion and, thus, the merging of the viral and host cell membranes in a pH-dependent fashion at the lysosomal membrane. The FD consists of two distinct regions: an N-terminal fusion peptide (FP, G^260^ – T^274^) and an internal fusion loop (FL, C^279^ – N^295^) that are connected by a short linker region (P^275^ – Y^278^). Nonetheless, the precise structural and functional characteristics of the LASV FD remain unknown. Here, we demonstrate that the LASV FD associates with the host cell membrane via its FL, specifically residues R^282^ – L^290^, whereas the FP is more solvent-exposed, especially for residues D^268^ – T^274^. We found that a multitude of conformational states are adopted by the entire LASV FD before membrane association, while only the FP, and not the FL, continues to sample numerous states after membrane association. Moreover, we provide evidence that the LASV FD prefers to interact with anionic lipids, namely bis(monoacylglycero)phosphate (BMP). In conclusion, our findings indicate that the LASV FD preferentially initiates fusion in the presence of BMP, at which point the FL adopts a helical conformation to associate with the membrane, whereas the FP remains exposed to the environment.

## Introduction

Lassa Virus (LASV), a member of the Arenaviridae family, is a zoonotic pathogen responsible for Lassa fever (LF) – a severe hemorrhagic fever that poses a significant public health threat, chiefly in West Africa where it is endemic^1–4^. Annually, an estimated 100,000 to 300,000 individuals are infected with LASV, but poor healthcare in endemic regions has caused an underrepresentation of cases. The case fatality rate (CFR) of LASV historically averages around 20%, especially in severe cases and pregnant individuals, leading to thousands of annual deaths^5–8^. LASV is most commonly spread by contact with excrement from infected *Mastomys natalensis* rodents, but an increased incidence of direct human-to-human transmission has been observed in recent years^9–11^. Despite its clinical importance, there are presently no FDA-approved therapeutic options for the explicit treatment of LF^12, 13^. It thus comes as no surprise that the World Health Organization (WHO) has categorized LASV as one of the top five infectious agents requiring prioritized research and development due to its pandemic potential, should it complete its zoonotic jump^14, 15^.

LASV relies on membrane fusion to enter the host cell, a process facilitated by the glycoprotein complex (GPC)^16–18^. The GPC is the sole protein on the virion surface and is comprised of a receptor binding subunit (glycoprotein 1, GP1) and a fusion subunit (glycoprotein 2, GP2) that is associated with a stable signal peptide (SSP). After an initial interaction between GP1 and its primary receptor, α-dystroglycan (αDG), which is located on the host cell plasma membrane, the virion is endocytosed and delivered late within the endocytic pathway where fusion ultimately occurs via the six-helix bundle mechanism (6HB)^19–23^. More specifically, the low pH environment triggers GP1 to undergo conformational changes, allowing it to engage with its secondary receptor, lysosomal-associated membrane protein 1 (LAMP1), and dissociate from the GPC^24–28^. A hydrophobic sequence at the N-terminus of GP2, known as the fusion domain (FD), is subsequently exposed and anchors itself into the host cell membrane. This results in the initiation membrane fusion, leading to the formation of a fusion pore and delivery of LASV’s genetic material into the host cell. Thus, it is evident that the LASV FD has an essential role in viral entry and its structure before (pre-) and after (post-) fusion should be further explored.

LASV is a class one fusion protein, similar to human immunodeficiency virus (HIV), influenza, Ebola virus (EBOV), and severe acute respiratory coronavirus 2 (SARS-CoV-2). Typically, the FD of class one fusion proteins is an N-terminal fusion peptide (FP), as seen in HIV and influenza^29–36^. In some cases, however, an internal fusion loop (FL) has been observed, such as that found in EBOV^37, 38^. The FD of LASV and other arenaviruses unusually contains both an FP (G^260^ – T^274^) and FL (C^279^ – N^295^), which are connected by a short linker region (P^275^ – Y^278^) [Figure 1]^39–42^. Notably, the internal disulfide bond within the FL is located between residues C^279^ and C^292^. This feature is shared only with coronaviruses, which includes SARS-CoV-2, the causative agent of coronavirus disease 2019 (COVID-19). While the SARS-CoV-2 FP and FL can function independently, they are most efficient in synergy^43–50^. It has been postulated that the SARS-CoV-2 FD functions as an evolutionary hybrid, potentially contributing to the remarkable infectivity of COVID-19. However, many aspects of the LASV FD’s molecular mechanisms, particularly those governing its ability to mediate fusion and the roles of both the FP and FL, remain poorly understood. In the literature, it has been demonstrated that the LASV FD undergoes a significant conformational change from the pre- to post-fusion state, as it transitions from a nonfusogenic coil to a fusogenic helix that can associate with the target host cell membrane, primarily the lysosomal membrane^41^. For clarity, the nonfusogenic coil conformation will be referred to as the pre-fusion state, and fusogenic helix conformation as the post-fusion state in this manuscript. Nonetheless, the exact location of the helix, residues that perturb the membrane, and influence of lipids found within the lysosomal membrane are unknown.

**Figure 1.**
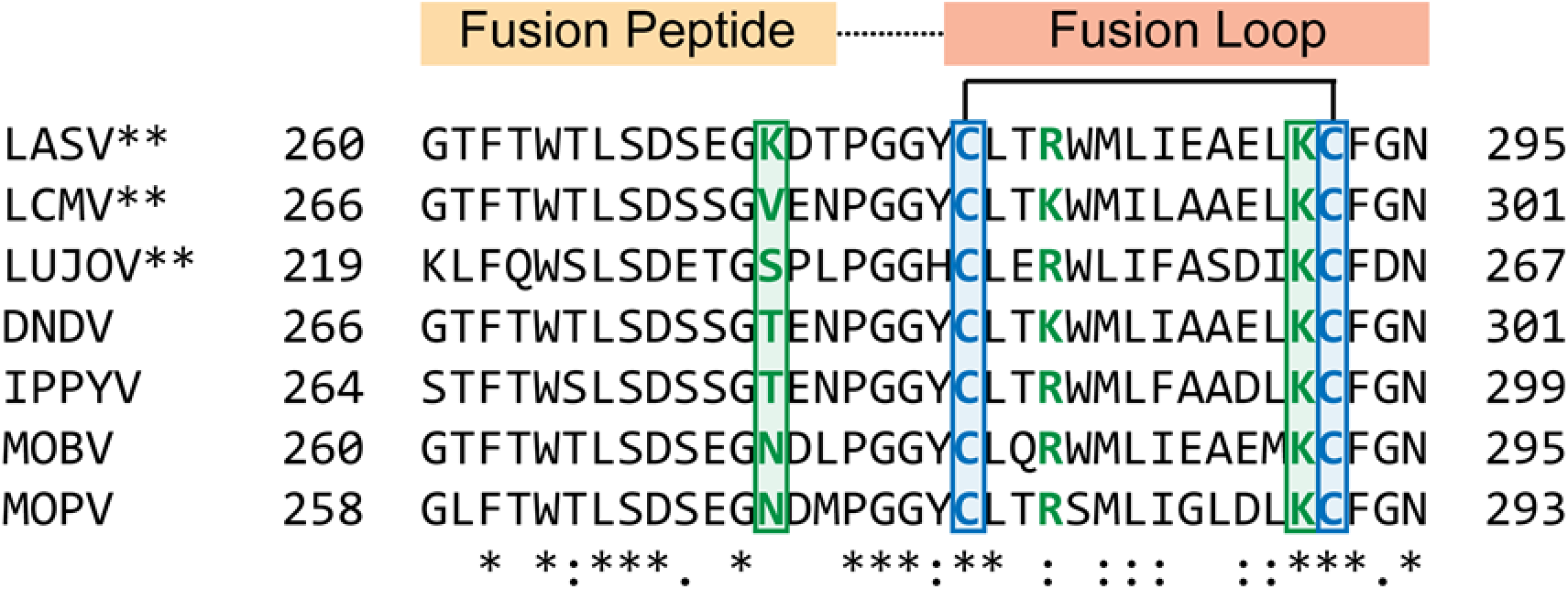
Multiple sequence alignment of the fusion domain (FD) in the Old-World clade of the arenavirus family. The structurally distinct regions that make up the FD, the fusion peptide (FP, yellow) and internal fusion loop (FL, orange), are conjoined by a short linker region. Positively charged residues within the LASV FD are highlighted (bold, green) with those studied in this article denoted (box). Viruses with pathogenic potential in humans are indicated by **. Residues with full conservation are indicated by an asterisk (*), whereas those with strong and weak similarities are denoted by a semicolon (:) and period (.), respectively. Abbreviations: LCMV, lymphocytic choriomeningitis virus; LUJOV, Lujo virus; DNDV, Dandenong virus; IPPYV, Ippy virus; MOBV, Mobala virus; MOPV, Mopeia virus.

In this study, we aimed to elucidate the structure of the LASV FD in its pre- and post-fusion states and the functional influence of key lipids found within the lysosomal membrane. Through a combination of NMR techniques, we illustrate a conformational change undergone by the LASV FD following its association with the membrane. In the post-fusion state, the FL, namely residues R^282^ –L^290^, adopts a helix that embeds itself just below the lipid head group of the membrane. In contrast, the FP, mainly residues D^268^ –T^274^, remains exposed to the aqueous environment. Multiple conformational states are adopted by both the FP and FL in the pre-fusion state, but this only applies to the FP after fusion occurs. Additionally, the LASV FD is partial to associate, and thus, initiate fusion, in membranes comprised of anionic lipids, especially bis(monoacylglycero)phosphate (BMP). We provide evidence that this interaction is not facilitated by positively charged, lysine residues (K^272^ and K^291^) as mutagenesis of these residues did not have significant impacts on secondary structure, membrane affinity, or fusion. Together, this study illuminates how the LASV FD associates with the membrane and shows its preference to initiate fusion with anionic lipids, chiefly BMP, which is not mediated by an ionic interaction with lysine residues.

## Materials And Methods

### Lipids

1,2-dimyristoyl-sn-glycero-3-[phospho-rac-(1-glycerol)] (DMPG) was purchased from Anatrace (Maumee, OH, USA). 16:0–18:1 1-palmitoyl-2-oleoyl-glycero-3-phosphocholine (POPC), 16:0–18:1 1-palmitoyl-2-oleoyl-*sn*-glycero-3-phospho-[1′-rac-glycerol] (POPG) [Figure S1A], 1,2-dioleoyl-*sn*-glycero-3-[phospho-rac-(3-lysyl(1-glycerol))] (DOPG) [Figure S1B], 1-palmitoyl-2-oleoyl-*sn*-glycero-3-phosphate (POPA) [Figure S1C], 1,2-dioleoyl-*sn*-glycero-3-phosphate (DOPA) [Figure S1D], 1-palmitoyl-2-oleoyl-*sn*-glycero-3-phospho-L-serine (POPS) [Figure S1E], 1,2-dioleoyl-*sn*-glycero-3-phospho-L-serine (DOPS) [Figure S1F], 1-palmitoyl-2-oleoyl-*sn*-glycero-3-phosphoethanolamine (POPE) [Figure S1G], bis(monooleoylglycero)phosphate (S,R Isomer) (BMP) [Figure S1H], 18:1 1,2-dioleoyl-*sn*-glycero-3-phosphoethanolamine-*N*-[lissamine rhodamine B sulfonyl] (Rh-PE), 18:1 1,2-dioleoyl-*sn*-glycero-3-phosphoethanolamine-*N*-[7-nitro-2–1,3-benzoxadiazol-4-yl] (NBD-PE), 1,2-dimyristoyl-*sn*-glycero-3-phosphocholine (DMPC), and 1,2-dihexanoyl-*sn*-glycero-3-phosphocholine (DHPC) were all purchased from Avanti Polar Lipids (Alabaster, AL, USA).

### Expression and Isotopic labeling

The expression protocol and single, ^15^N-labeling of the Lassa virus fusion domain (LASV FD) has been previously described^41, 51^. The lysine mutants (i.e., K^272^A and K^291^A) followed the designed expression protocol. However, the F^293^W-containing vector was transformed into *Escherichia coli* strain DL39(DE3) auxotrophic cells instead of *E. coli* strain BL21(DE3)pLysS cells. For selective, ^19^F-labeling, a single colony from the transformation was used to inoculate 5 mL of LB media containing 50 μg/mL kanamycin (ThermoFisher Scientific; Waltham, MA, USA) and grown overnight (16 – 18 h) at 37 ℃ and 225 rpm in a MaxQ 4000 incubated shaker (ThermoFisher Scientific; Waltham, MA, USA). The next morning, the starter culture was added to 1 L of MDAG minimal media (25 mM Na_2_HPO_4_, 25 mM KH_2_PO_4_, 50 mM NH_4_Cl, 5 mM Na_2_SO_4_, 2 mM MgSO_4_, 0.2X trace metals, 0.5% glucose, 50 μg/mL kanamycin, pH 7.2) supplemented with 200 μg/mL of each amino acid, except for Cys and Tyr, which were not added due to solubility issues^52^. The trace metals came from a 1000X stock that contained 50 mM FeCl_3_, 20 mM CaCl_2_, 10 mM MnCl_2_, 10 mM ZnSO_4_, and 2 mM each of CoCl_2_, CuCl_2_, NiCl_2_, Na_2_MoO_4_, Na_2_SeO_3_, and H_3_BO_3_ in 60 mM HCl. The cells were grown at 37 ℃ and 225 rpm until an OD_600_ of 0.6 was achieved, as confirmed on an Ultrospec 1000 Spectrophotometer (Pharmacia Biotech; Cambridge, England, GBR), then harvested for 20 min at 4,000 x g and 4 ℃ in an Avanti J-15R centrifuge (Beckman Coulter; Brea, CA, USA). Each pellet was subsequently resuspended in 250 mL of MDAG minimal media that lacked any amino acids (25 mM Na_2_HPO_4_, 25 mM KH_2_PO_4_, 50 mM NH_4_Cl, 5 mM Na_2_SO_4_, 2 mM MgSO_4_, 0.2X trace metals, 0.5% glucose, 50 μg/mL kanamycin, pH 7.2), then centrifuged for 20 min at 4,000 x g and 4 ℃ in an Avanti J-15R centrifuge (Beckman Coulter; Brea, CA, USA) to remove any residual natural phenylalanine from the system. The rinsed cell pellets were resuspended in 1 L of ^19^F-Phe MDAG minimal media (100 mg 4-fluoro-D,L-phenylalanine (Fisher Scientific; Hampton, NH, USA), 25 mM Na_2_HPO_4_, 25 mM KH_2_PO_4_, 50 mM NH_4_Cl, 5 mM Na_2_SO_4_, 2 mM MgSO_4_, 0.2X trace metals, 0.5% glucose, 50 μg/mL kanamycin, pH 7.2) with 200 μg/mL of each amino acid except Cys, Try, and Phe. The resuspended cells were shaken for 30 min at 37 ℃ and 225 rpm in a MaxQ 4000 incubated shaker (ThermoFisher Scientific; Waltham, MA, USA) to give the cells time to adjust to the new media. After 30 min, the temperature was decreased to 18 ℃ and the cells were shaken at 225 rpm for an additional 30 min in a MaxQ 4000 incubated shaker (ThermoFisher Scientific; Waltham, MA, USA) to allow the cells to acclimate to the new temperature. Protein expression was subsequently induced with 1 mM isopropyl β-D-1-thiogalactopyranoside (IPTG), and then the cells were incubated overnight (∼20 h) before being harvested at 4,000 x g and 4 ℃ for 45 min in an Avanti J-15R centrifuge (Beckman Coulter; Brea, CA, USA).

Selective, ^15^N-Leu labeling of the LASV FD was achieved in a similar manner as site specific, ^19^F labeling with some modifications. Initially, 4 x 120 mL propagated starter cultures of the LASV FD in *E. coli* BL21(de3)pLysS cells were added to 4 x 1 L of EMBL minimal media (1 g/L NH_4_Cl, 10 g/L glucose, 8 g/L Na_2_HPO_4_, 2 g/L KH_2_PO_4_, 0.5 g/L NaCl, 1 mM MgSO_4_, 0.3 mM Na_2_SO_4_, 0.3 mM CaCl_2_, 50 μg/mL kanamycin, 34 μg/mL chloramphenicol, trace amounts of biotin and thiamine) at 37 ℃ and 225 rpm in a MaxQ 4000 incubated shaker (ThermoFisher Scientific; Waltham, MA, USA) until an OD_600_ between 0.6 and 0.8 was achieved, as measured on an Ultrospec 1000 Spectrophotometer (Pharmacia Biotech; Cambridge, England, GBR). Upon reaching the appropriate OD_600_ , cells were harvested for 15 min at 4,000 x g and 15 ℃ in an Avanti J-15R centrifuge (Beckman Coulter; Brea, CA, USA) such that each pellet contained 1 L worth of cells. Each pellet was subsequently resuspended in 250 mL of EMBL minimal media that lacked a nitrogen source (10 g/L glucose, 8 g/L Na_2_HPO_4_, 2 g/L KH_2_PO_4_, 0.5 g/L NaCl, 1 mM MgSO_4_, 0.3 mM Na_2_SO_4_, 0.3 mM CaCl_2_, 50 μg/mL kanamycin, 34 μg/mL chloramphenicol, trace amounts of biotin and thiamine) then spun down for 15 min at 4,000 x g and 15 ℃ to remove any residual unlabeled media from the system. The rinsed cell pellets were resuspended in 1 L of ^15^N-Leu EMBL minimal media (1 g/L ^15^N-Leu (Cambridge Isotope Laboratories; Tewksbury, MA, USA), 10 g/L glucose, 8 g/L Na_2_HPO_4_, 2 g/L KH_2_PO_4_, 0.5 g/L NaCl, 1 mM MgSO_4_, 0.3 mM Na_2_SO_4_, 0.3 mM CaCl_2_, 50 μg/mL kanamycin, 34 μg/mL chloramphenicol, trace amounts of biotin and thiamine) with 10 g of each, unlabeled amino acid, except for Leu and Tyr. The addition of unlabeled amino acids in 10-fold excess over the desired amino acid to incorporate (Leu) reduces scrambling during protein expression, whereas Tyr was not included due to solubility issues^52, 53^. The resuspended cells were split into 2 x 500 mL and grown for 1 h at 37 ℃ and 225 rpm in a MaxQ 4000 incubated shaker (ThermoFisher Scientific; Waltham, MA, USA) to acclimate the cells to the new media and temperature. Cells were then induced with 1 mM isopropyl β-D-1-thiogalactopyranoside (IPTG) and incubated overnight (∼20 h) before being harvested at 4,000 x g and 4 ℃ for 45 min in an Avanti J-15R centrifuge (Beckman Coulter; Brea, CA, USA).

For double, ^15^N/^13^C isotopic labeling of the LASV FD, a comparable method to ^15^N-Leu labeling was employed. However, cells were initially grown in 4 x 1 L of LB until the appropriate OD_600_, rinsed with 250 mL of EMBL minimal media lacking a carbon or nitrogen source (8 g/L Na_2_HPO_4_, 2 g/L KH_2_PO_4_, 0.5 g/L NaCl, 1 mM MgSO_4_, 0.3 mM Na_2_SO_4_, 0.3 mM CaCl_2_, 50 μg/mL kanamycin, 34 μg/mL chloramphenicol, trace amounts of biotin and thiamine), then resuspended in 1 L of ^15^N/^13^C EMBL minimal media (1 g/L ^15^NH_4_Cl (Cambridge Isotope Laboratories; Tewksbury, MA, USA), 2 g/L ^13^C glucose (Cambridge Isotope Laboratories; Tewksbury, MA, USA), 8 g/L Na_2_HPO_4_, 2 g/L KH_2_PO_4_, 0.5 g/L NaCl, 1 mM MgSO_4_, 0.3 mM Na_2_SO_4_, 0.3 mM CaCl_2_, 50 μg/mL kanamycin, 34 μg/mL chloramphenicol, trace amounts of biotin and thiamine). The cells were grown for1 h at 37 ℃ and 225 rpm in a MaxQ 4000 incubated shaker (ThermoFisher Scientific; Waltham, MA, USA) before being induced with 1 mM IPTG, incubated for an additional 4 h at 37 ℃ and 225 rpm in a MaxQ 4000 incubated shaker (ThermoFisher Scientific; Waltham, MA, USA), and then harvested at 4,000 x g and 4 ℃ for 45 min in an Avanti J-15R centrifuge (Beckman Coulter; Brea, CA, USA). For triple, ^2^H/^15^N/^13^C isotopic labeling, 1L of D_2_O was utilized to prepare the ^15^N/^13^C EMBL minimal media as opposed to H_2_O but expressed in the same manner as the ^15^N/^13^C sample; however, after induction with 1 mM IPTG, the cells were incubated for 8 h at 37 ℃ and 225 rpm, then harvested with the same parameters. Cell pellets were stored at -80 ℃ or immediately purified.

### Purification

The purification protocol of the Lassa virus fusion domain (LASV FD) has been described in detail previously^41, 51^. Briefly, a construct containing the entire FD [^260^(GTFTWTLSDSEGKDTPGGYCLTRWMLIEAELKCFGN)^295^] was designed with an N-terminal 9x-His tag followed by a trp operon leader sequence (TrpLE), and thrombin cleavage site. The natural thrombin cleavage site was replaced with LVPR↓GT to yield the native sequence of the LASV FD after cleavage. Ni-NTA affinity chromatography and thrombin cleavage were employed to separate the protein from the tags. The separated protein was then subjected to size exclusion chromatography (SEC) with HMA buffer (10 mM HEPES/MES/NaOAc, 100 mM NaCl, pH 7.4) or NMR buffer (25 mM Na_2_HPO_4_, 100 mM NaCl, pH 7.0) to further isolate the protein and achieve the correct buffer system. All mutants (i.e., K^272^A, K^291^A, and F^293^W) in this study were otherwise purified in the same manner as the LASV FD (WT).

### Nuclear Magnetic Resonance (NMR) experiments

NMR spectra were acquired utilizing a Shigemi NMR tube with a sample volume of ∼300 μL in NMR buffer, pH 7.0 (pre-fusion) or pH 4.0 (post-fusion) and 9:1 ratio of H_2_O:D_2_O. The protein concentration was ∼500 μM for single, ^15^N labeled samples, ∼650 μM for double, ^15^N/^13^C labeled samples, ∼1,000 μM for the ^15^N-Leu labeled sample, and 1,000 μM for the ^19^F labeled samples. For the post-fusion state, 25% acidic bicelles ((75:25 DMPC:DMPG):DHPC) with a q-value of 0.5 were combined with the sample, then the pH was dropped to 4.0 with 1 M HCl and verified with a calibrated pH probe. The q-value was confirmed via phosphorous experiments on an Ascend 800 MHz magnet (Bruker; Billerica, MA, USA) equipped with a CPQCI ^1^H-^31^P/^13^C/^15^N/D Z-GRD Cryoprobe. Experiments were carried out on either the Ascend 800 MHz magnet or an Ultrashield 600 MHz magnet (Bruker; Billerica, MA, USA) with a CP2.1 TCI 600S3 H&F/C/N-D-05 Z XT Cryoprobe at a temperature of either 20 ℃ (pre-fusion) or 45 ℃ (post-fusion) to improve peak sharpness [Figure S2]. A backbone assignment was completed for both states following HNCA, HN(CO)CA, HNCO, HN(CA)CO, and HN(CA)CB experiments using either 20% or 25% non-uniform sampling schedule (NUS). An HN(CA)CB experiment was run on the triple-labeled (^2^H/^15^N/^13^C) sample for the pre-fusion state, whereas HNCA, HNCO, HN(CA)CO, and HN(CA)CB experiments were run for the post-fusion state to improve signal-to-noise for the backbone assignment. All backbone experiments were processed via NMRPipe^54^ and NMRFAM-SPARKY^55^ via NMRBox^56^. The chemical shift index (CSI) for each residue was deduced from the equation CSI = Cα_FD_ – Cα_BMRB_ where Cα_FD_ is the Cα measured for the LASV FD and Cα_BMRB_ is the Cα of a given residue as published in the Biological Molecular Resonance Bank (BMRB)^57, 58^. CSI of N, H, and CO were performed in the same manner but with the value of N, H, or CO from BMRB subtracted from the N/H/CO of the FD. A chemical shift list was exported from NMRFAM-SPARKY in a format suitable for Torsion Angle Likelihood Obtained from Shift and Sequence Similarity (TALOS+) with the tf command^55, 59^. The file was adjusted to have the input table with the required data format, then the TALOS+ prediction was performed with the -iso option in the command line to correct the ^13^Cα and ^13^Cβ due to ^2^H isotopic labeling. Visual analysis of the resultant secondary structure was conducted on a Ramachandran map. Integrations for the multiple conformers of each residue were calculated in Bruker TopSpin, and then the percentage of each conformer was calculated from 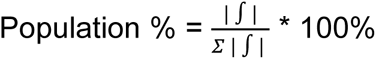 where |∫ | is the absolute integration of a conformer and Σ | ∫ | is the sum of the absolute integration for all conformers of a given residue. The Poky Suite was used to extract ^1^H slices from the ^1^H – ^15^N heteronuclear single quantum coherence (HSQC) of the pre- and post-fusion state to visualize the populations of the different conformers for a given residue^60^. All ^19^F experiments were carried out on the Ultrashield 600 MHz magnet (Bruker; Billerica, MA, USA) with a CP2.1 TCI 600S3 H&F/C/N-D-05 Z XT Cryoprobe, then processed using the software MestRe Nova (Mnova) within NMRBox.

### Dynamic experiments

Heteronuclear (^1^H – ^15^N) NOEs, T_1_, and T_2_ experiments at pH 7.0 and pH 4.0 were conducted on the Ascend™ 800 MHz magnet (Bruker; Billerica, MA, USA) equipped with a CPQCI ^1^H-^31^P/^13^C/^15^N/D Z-GRD Cryoprobe. Spin-lattice T_1_ relaxation experiments were performed with a variable delay (vd) list of either 0.1, 0.3, 0.5, 0.8, 1.5, and 3 s or 0.1, 0.2, 0.5, 0.8, 1, 2, and 4 s for the pre- and post-fusion states, respectively. Spin-spin T_2_ relaxation experiments had a variable counter (vc) list of either 2, 4, 6, 8, 10, 20, 30, and 40 or 2, 4, 6, 8, 10, 15, 20, 30, and 40 for the pre- and post-fusion states, correspondingly, yielding relaxation times of 0.03392, 0.06784, 0.10176, 0.13568, 0.1696, 0.2544, 0.5088, and 0.6784 s or 0.03392, 0.06784 0.10176, 0.13568, 0.1696, 0.2544, 0.3392, 0.5088 and 0.6784 s for the loops accordingly. Heteronuclear NOEs were processed via NMRPipe^54^ and NMRFAM-SPARKY^55^ via NMRBox^56^. Data from the T_1_ and T_2_ experiments was processed in the Bruker TopSpin Dynamics Center, then converted to R_1_ and R_2_ by taking the inverse of T_1_ and T_2_, correspondingly. The error shown is propagated from the signal-to-noise or standard error of the mean (SEM).

### Paramagnetic Relaxation Enhancement (PRE) experiments

Membrane depth was probed utilizing gadolinium (III) diethylenetriaminepentaacetic acid (Gd-DTPA), 5-doxyl stearic acid (5-DSA), and 16-doxyl stearic acid (16-DSA). 100 mM Gd-DTPA was solubilized in H_2_O and then titrated into the NMR sample to achieve 0.5, 1, 2, 4, and 6 mM. For 5- and 16-DSA, known concentrations of DSA were added to glass tubes. A film was created by gently vortexing the tube under a nitrogen stream to remove the chloroform. The NMR sample was subsequently added to the glass tube to resuspend the DSA film and incorporate the paramagnetically tagged stearic acid into the system. The 5- and 16-DSA titrations were carried out to achieve concentrations of 0.5, 1, 2, 4, 6, and 8 mM. All data were collected through a ^1^H-^15^N HSQCat either 20 ℃ (pre-fusion) or 45 ℃ (post-fusion). All spectra were processed using NMRPipe and NMRFAM-SPARKY via NMRBox to obtain peak intensities and signal-to-noise ratios for the titration of each relaxation agent. Relative intensities were calculated as 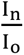 where I_n_ is the peak intensity at titration point n and I_o_ is the maximum peak intensity. The error shown is from the signal-to-noise ratio or SEM.

### Preparation of Unilamellar Vesicles

Large unilamellar vesicles (LUVs) were prepared by combining specified amounts of lipid stock solutions in glass test tubes. Chloroform was removed by applying a continuous nitrogen stream while gently vortexing the glass tube to create a lipid film before being left in a vacuum desiccator overnight to evaporate any residual chloroform. The following day, the lipid film was resuspended via vortexing in the appropriate volume of HMA buffer, subjected to 10 freeze-thaw cycles between liquid nitrogen and a 42 ℃ water bath, then extruded 21 times through a double layer of 100 nm pore-size polycarbonate membranes (Avestin; Ottawa, ON, CAN). The lipid film for small unilamellar vesicles (SUVs) was created analogous to the LUVs but was resuspended in the appropriate volume of either CD buffer (1 mM HMA, 10 mM NaCl, pH 7.4) or ITC buffer (10 mM NaOAc, 100 mM NaCl, pH 4.0), then sonicated on ice for 15 total min (1 s on, 1 s off, 10% power) using a Branson® ultrasonicator furnished with a titanium microtip (Emerson; Danbury, CT, USA). The transparent solution was subsequently centrifuged for 15 min at 20,000 x g in an Eppendorf 5425 microcentrifuge (Sigma-Aldrich; St. Louis, MO, USA)to remove any particulates and transferred to a new microcentrifuge tube. All vesicles used in this study were comprised of 65 mol% POPC:35 mol% POPG, unless otherwise specified.

### FRET-Based Fusion Assay

Unlabeled LUVs comprised of 65:35 POPC:POPG and labeled LUVs of 63:35:1:1 POPC:POPG:Rh-PE:NBD-PE were mixed at a ratio of 9:1 unlabeled:labeled, unless otherwise indicated. 5 μM protein, whether that be the WT or each mutant, and 100 μM LUVs were mixed to achieve a protein:lipid ratio of 1:20 in HMA buffer. All experiments were carried out in a Corning Costar black-walled, clear-bottomed 96-well plate with 150 μL per well. Fluorescence was recorded at room temperature (∼23℃) on a SpectraMax M5 microplate reader (Molecular Devices; San Jose, CA, USA) with excitation and emission wavelengths recorded at 460 nm and 538 nm, accordingly, with a cutoff at 530 nm. Percent fusion was calculated from the equation 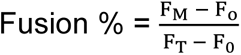 *100% where F_M_ is the measured fluorescence at lysosomal pH (4.0), F_o_ is the background fluorescence at physiological pH (7.4), and F_T_ is the total fluorescence after 1% Triton X-100 was added to the system to cause complete vesicle disruption. Acidification was carefully controlled through the addition of 1 M HCl, as verified by a pH titration of the HMA buffer, which is linear from pH 3.0 – 8.0. Controls containing no protein, merely LUVs and HMA buffer, were run alongside each experimental condition, subjected to the same equation, and subtracted from the final values with error propagated from the SEM.

### Circular Dichroism (CD) Spectroscopy

To prepare the protein, either WT or each mutant, for CD experiments, the sample was diluted with 15 mL of CD buffer, then concentrated down to 1 mL with a 15 mL capacity, 3 kDa MWCO Amicon Ultra Centrifugal Filter (Sigma-Aldrich; St. Louis, MO, USA). This was repeated five times to fully exchange the sample from HMA buffer to CD buffer, where the latter is 10x diluted to prevent noisy signals from excess salt concentrations. CD measurements were conducted at room temperature (∼23 ℃) on a Jasco J-810 spectrometer (Jasco; Easton, MD, USA) utilizing a quartz cuvette with a 2 mm pathlength. Samples were comprised of 8 μM of WT or each mutant and 0.8 mM SUVs in CD buffer for a ratio of 1:100 protein:lipid. Spectra were collected from 260 nm to 198 nm with a step size of 1 nm at 20 nm/min and averaged over three accumulations. Control measurements were taken for 0.8 mM SUVs alone in CD buffer. CDToolX was employed to subtract the control measurements from the respective sample, then the spectra were smoothed via the software^61^. Spectra were converted to mean residue ellipticity (MRE) units. Helical percentages were calculated from the MRE of a given spectrum at 222 nm (θ_222_) from the equation 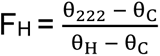 where θ_C_ = 2,220 – 53T, θ_H_ = (250T – 44,000)*(1 - 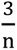), T is the temperature in Celsius, and n is the number of residues in the protein^37, 41, 62, 63^.

### Isothermal Titration Calorimetry (ITC)

The desired protein, either WT or each mutant, was added to a 0.5 – 3 mL, 3.5 kDa MWCO Slide-A-Lyzer™ dialysis cassette (Thermo Fisher Scientific; Waltham, MA, USA) and dialyzed against 2 L of ITC buffer at 4 ℃ for a minimum of 4 h. The protein was subsequently removed from the dialysis cassette, spun down for 15 min at 20,000 x g in an Eppendorf 5425 microcentrifuge (Sigma-Aldrich; St. Louis, MO, USA), and transferred to a clean microcentrifuge tube. The protein was then degassed for 8 min at room temperature in a Microcal Thermovac alongside 20 mM of 65:35 POPC:POPG SUVs that were in the ITC buffer. The microplate reader protocol for the Pierce™ Bicinchoninic Acid (BCA) protein assay kit (Thermo Fisher Scientific; Waltham, MA, USA) was employed to determine the protein concentration of the resulting sample, which was typically ∼15 – 20 μM, then ran on the ITC. A Malvern VP-ITC microcalorimeter (Malvern Panalytical; Malvern, England, GBR) was used for all ITC measurements where the sample cell was filled with the degassed protein and the injection syringe with the degassed SUVs. A total of 41 injections were performed at 23 ℃ with the reference power set to 5 μcal/s, stirring speed of 310 rpm, and filter period of 2 s. Each injection was set to 7 μL with a 16.8 s duration, except for the first injection which was 2 μL with a 4.8 s duration. Al processing was conducted through NITPIC^64^ and SEDPHAT^65^ with final figures created via GUSSI^66^. At least two titrations were performed for each condition to deduce the average K_d_ and standard deviation, which are presented on each isotherm.

### Molecular Dynamics (MD) Simulations

The initial structural model of the LASV FD was taken from the Protein Data Bank (PDB) entry 7PUY^16^ using residues G^260^ – N^295^. The terminal amino acids were capped using the acetylated N-terminal (ACE) and methyl-amidated C-terminal (CT3) blocking groups^49, 67, 68^. A series of membrane systems composed of POPC and either POPG, POPS, or BMP were built using CHARMM-GUI *Membrane Builder* [Table S1]^69–71^. All simulations were performed with all-atom CHARMM36(m) force fields for the protein and lipids^72, 73^, whereas TIP3P parameters were used for water with sodium and chloride ions added to reach a neutral concentration of 150 mM NaCl^74^. The pH of the system was set to 4.0 with all aspartic acid (D^268^) and glutamic acid (E^270^, E^287^, and E^289^) residues protonated. The temperature and pressure of the system were set to 318.15 K and 1 bar, accordingly. The protonated LASV FD was initially translated approximately 20 Å above the membrane. The initial system size of each replica was 100 Å × 100 Å x 128 Å to accommodate all initial components, yielding a total of roughly 130,000 atoms per replica. 2 μs molecular dynamics (MD) simulations were performed for five replicas for each of the three membrane systems, leading to a total of fifteen independent simulations. All simulations were conducted using OpenMM^75^ with the inputs generated by Membrane Builder^76, 77^. Analyses were performed using MDanalysis^78, 79^ and models were visualized in VMD, version 1.9.4^80^.

## Results

### A conformational change of the LASV FD at a low pH results in its FL adopting a helix and associating with the host cell membrane

We utilized a series of solution NMR spectroscopy backbone experiments to investigate the structure of the LASV FD in both its pre- and post-fusion states. For the pre-fusion state, G^260^ and T^261^ could not be assigned, due to their inherent flexibility as terminal residues, whereas I^286^ of the post-fusion backbone was not assigned due to peak ambiguity, correlating with 95% pre- and 97% post-fusion backbone assignment rates, respectively. A comparison of ^1^H – ^15^N heteronuclear single quantum coherence (HSQC) spectra of the FD in solution at physiological pH [Figure 2A] and with acidic bicelles at lysosomal pH [Figure 2B] revealed moderate chemical shift perturbations, suggestive of a conformational change. To rudimentarily estimate the secondary structure of the LASV FD, we utilized the backbone assignment for the pre- and post-fusion states to generate a chemical shift index (CSI). The chemical shift of the alpha carbons (Cα) for all residues within the FD was compared to that of a random coil where a positive value greater than one indicates a helix, a negative value indicates a sheet, and a value close to zero indicates a random coil^58^. The CSI suggests an almost completely random coil secondary structure for both the pre- and post-fusion states [Figure S3A]. The same trend was observed for the CSI of the amide (N) [Figure S3B], amide hydrogen (H) [Figure S3C], and carbonyl carbons (CO) [Figure S3D]. While the CSI of the pre-fusion state agreed with the literature^16, 17, 41^, the lack of helical structure in the post-fusion state was unexpected. Subsequently, we employed Torsion Angle Likelihood Obtained from Shift and Sequence Similarity (TALOS+) to yield a more in-depth secondary structural analysis of the LASV FD from our backbone assignment and provide predictions for the missing peaks of both states. This analysis further supported the notion that the pre-fusion state of the LASV FD is a random coil [Figure 2C (red); Table S2]. In the post-fusion, however, it was revealed that an alpha helix of approximately three turns exists within the FL for residues R^282^ – L^290^ [Figure 2C (blue); Table S3]. Additionally, TALOS+ indicated a short helix potentially formed for residues S^267^ – E^270^, but this would be unlikely due to the high propensity of destabilizing amino acids (i.e., S and D) and lack of hydrophobic amino acids. This corresponds well with the literature where it has been shown that the LASV FD adopts a helical structure with ∼13 residues involved in the post-fusion state, as previously shown via CD spectroscopy^41^. Notably, the internal disulfide bond is juxtaposed to this potential helix and formed correctly in both states. This is evidenced by the beta carbon (Cβ) chemical shifts being greater than 35 ppm [Figure S4] in both states, indicating oxidized cysteines, and a lack of free thiols under native conditions [Figure S5]. Given its proximity to the helix, the disulfide bond potentially restricts the length of the helix in the post-fusion state. Altogether, it is indicated that the LASV FD undergoes a conformational change from a random coil in the pre-fusion state to a helix in the post-fusion state, which forms for residues R^282^ – L^290^ within the FL and can thus associate with the host cell membrane during fusion.

**Figure 2.**
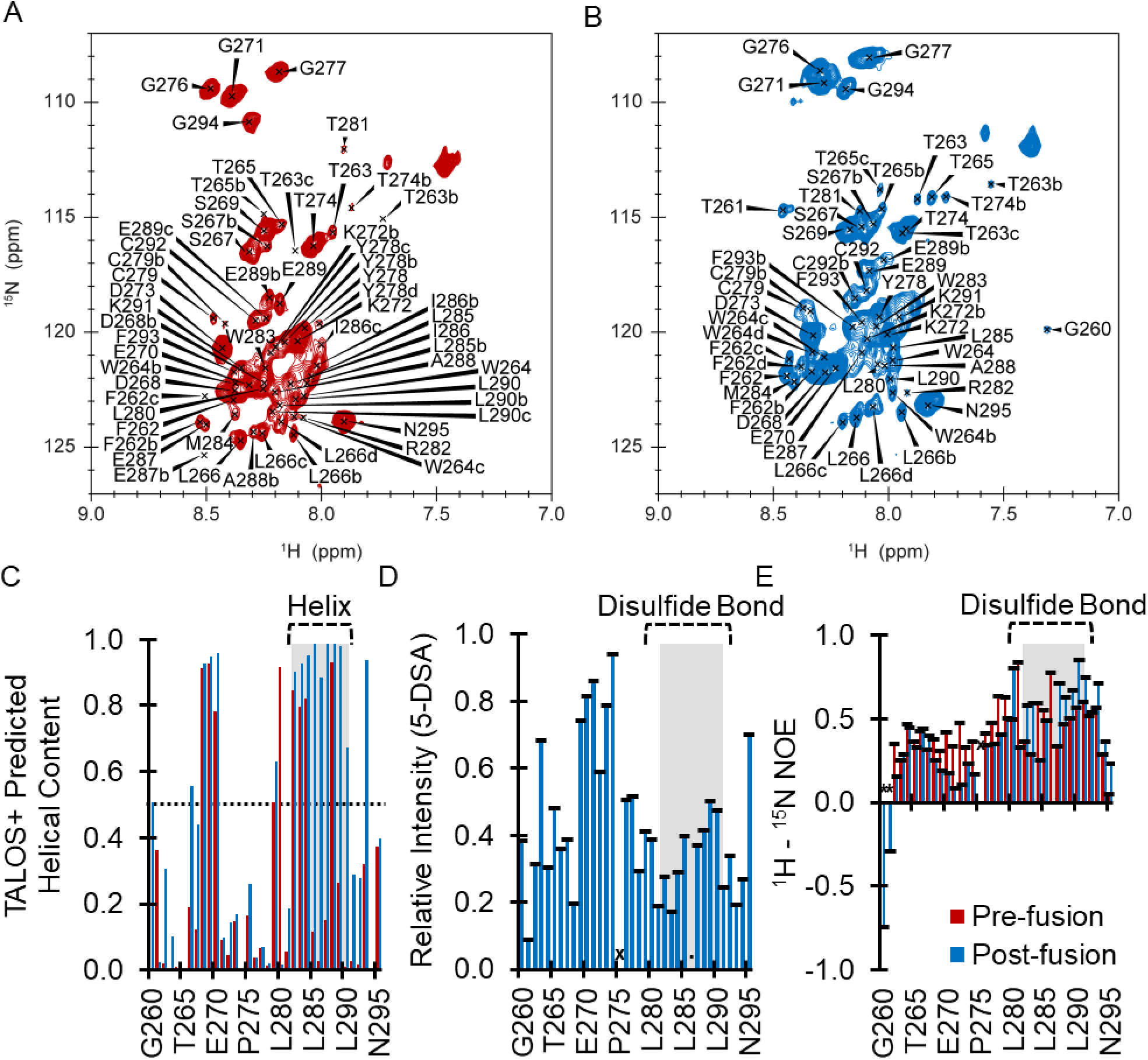
The LASV FD undergoes a conformational change from the pre- to post-fusion state and associates with the membrane via its FL. [A,B] ^1^H - ^15^N HSQC spectra highlight a multitude of peaks that shift from the [A] pre- to [B] post-fusion state. [C] An in-depth secondary structure analysis from the pre- (red) and post-fusion (blue) backbone assignment via TALOS+ provides predicted dihedral angles and suggests a helix is formed in the FL from R^282^ - L^290^ as values are consistently above 0.5. [D] Addition of 8 mM 5-DSA into the system containing the LASV FD resulted in quenching of the FL, but not the FP. [E] ^1^H - ^15^N NOE indicates that the FD becomes more restricted from the pre- (red) to post-fusion (blue) state with the FL being particularly limited. All measurements were carried out on ∼500 µM of the LASV FD in 300 µL of 25 mM Na_2_HPO_4_, 100 mM NaCl, pH 7.0 at 20 °c (pre-fusion) or pH 4.0 with acidic bicelles, q = 0.5 at 45 °c (post-fusion). Praline residues and other residues that could not be assigned for both states are marked with an X, whereas residues that could not be assigned for the pre- or post-fusion state are marked with an asterisk (*) or period (.), accordingly. The most populated conformer (a) is the residue label, whereas the second (b), third (c), and fourth (d) most populated conformers are shown. The disulfide bond (C^279^ and C^292^, dashed line) and helix (R^282^ - L^290^, transparent grey box) are indicated.

To further elucidate the depth of the LASV FD in the pre- and post-fusion states, we utilized paramagnetic relaxation enhancement (PRE) experiments with the paramagnetic probes Gd-DTPA, 5-DSA, and 16-DSA. The water-soluble Gd-DTPA causes signal quenching of solvent-exposed residues, whereas 5-DSA and 16-DSA quench signals for residues located proximally to the head and tail groups, respectively. In the pre-fusion state, the entire LASV FD (0.420 ± 0.021) experienced virtually the same amount of quenching as Gd-DTPA was titrated into the solution [Figure S5, Table S4]. The maximum quenching achieved for the LASV FP (0.458 ± 0.027) and FL (0.431 ± 0.025) were nearly identical, indicative that both were solvent-exposed. In the post-fusion state, however, we found that the FP was solvent-exposed, whereas the FL was inserted into the lipid head group of the membrane. The overall quenching of the FD in its post-fusion state (Gd-DTPA = 0.222 ± 0.026, 5-DSA = 0.439 ± 0.037, and 16-DSA = 0.620 ± 0.027) suggests that it associates with the membrane in a shallow manner. To be more specific, the relative intensities from the Gd-DTPA titration were 0.126 ± 0.022 and 0.317 ± 0.038 for the FP and FL, accordingly [Figure S7A]. A reverse trend was witnessed for the 5-DSA titration where the relative intensities of the FP were increased (0.530 ± 0.068) in comparison to the FL (0.353 ± 0.034) [Figure 2C]. Furthermore, residues D^268^ – T^274^ within the FP were significantly quenched by Gd-DTPA, but not 5-DSA, whereas residues R^282^ – L^285^ and K^291^ – F^293^ within the FL were quenched by 5-DSA, but not Gd-DTPA. The relative intensities from the titration of 16-DSA were nearly identical for the FP (0.623 ± 0.046) and the FL (0.620 ± 0.039) [Figure S7B]. Taking all of the PRE data into consideration, we believe that the FD associates with the membrane in a relatively shallow manner. The FL forms a boat-like helical structure for residues R^282^ – L^290^ that is embedded just below the lipid head groups. In contrast, the FP, mainly D^268^ – T^274^, does not insert itself into the membrane and is more solvent-exposed in both states.

We also investigated the protein dynamics of the LASV FD in the pre- and post-fusion states using ^1^H – ^15^N NOEs and R_1_ and R_2_ relaxation rates [Table S5]. The overall relaxation times of the FD in its pre-fusion state (^1^H – ^15^N NOEs = 0.448 ± 0.027 s [Figure 2E, red], R_1_ = 1.735 ± 0.091 s^-1^ [Figure S8A, red], and R_2_ = 4.941 ± 0.356 s^-1^ [Figure S8B, red]) agree relatively well with the dynamics of a random coil^45, 81^. In particular, the FP was slightly more flexible than the FL in the pre-fusion state, as denoted by the ^1^H – ^15^N NOEs of 0.369 ± 0.018 s and 0.497 ± 0.046 s, accordingly. In turn, for our relaxation experiments, we found that the FP had similar R_1_ relaxation times (1.750 ± 0.159 s^-1^) to the FL (1.651 ± 0.062 s^-1^), but lower T_2_ relaxation times (4.137 ± 0.550 s^-1^) than the FL (5.708 ± 0.501 s^-1^), further suggesting that the FP is somewhat more dynamic than the FL in the pre-fusion state. Opposingly, the relaxation times of the FD in its post-fusion state (^1^H – ^15^N NOEs = 0.419 ± 0.038 s [Figure 2E, blue], R_1_ = 1.305 ± 0.040 s^-1^[Figure S8A, blue], and R_2_ = 6.109 ± 0.683 s^-1^ [Figure S8B, blue]) are suggestive of a more restricted structural conformation. In the post-fusion state, the FP had much lower ^1^H – ^15^N NOEs (0.253 ± 0.034 s), particularly residues D^268^ – T^274^ than the FL (0.564 ± 0.048 s), indicating that the FP continued to be more flexible than the FL, but to a greater extent than in the pre-fusion state. Interestingly, the FP again had similar R_1_ relaxation times (1.280 ± 0.036 s^-1^) to the FL (1.339 ± 0.078 s^-1^), but lower R_2_ relaxation times (4.426 ± 0.390 s^-1^) than the FL (7.947 ± 1.224 s^-1^), further correlating with the FP being more dynamic than the FL in the post-fusion state. Taken together, these findings suggest that the two regions of the LASV FD have different dynamic properties in both states. While both are relatively flexible in the pre-fusion state, the FL becomes especially restricted in the post-fusion state due to its association with the membrane via residues R^282^ – L^290^, which are in a helical conformation, whereas the FP becomes more dynamic, chiefly for residues D^268^ – T^274^, likely due to its increased exposure to the solvent.

### Multiple conformations are adopted by certain residues in the pre- and post-fusion states of the LASV FD, as evidenced by a particularly dynamic leucine residue

When performing the backbone assignment, we noticed that multiple residues within the FD appeared to slowly sample several conformations in both the pre- and post-fusion states. We observed numerous ^1^H – ^15^N strips with the same chemical shifts for the Cα, Cβ, and CO, but different hydrogen and nitrogen shifts, of the particular residue (*i*). Additionally, in all of these strips, the chemical shifts of the Cα, Cβ, and CO were identical for the preceding residue (*i – 1*) as well. This suggested that a single residue was slowly exchanging between different conformations on a timescale that could be captured by the NMR as distinctive peaks, i.e., the millisecond to second scale^82–84^. The first residue was what we believed to be residue L^266^ located in the middle of the FP, which appeared to have distinct conformations in both the pre- and post-fusion states. In particular, we observed four different ^1^H – ^15^N strips with a Cα*_i_* of ∼55 ppm and Cβ*_i_* of ∼42 ppm, a trademark of leucine residues, and Cα*_i–1_* of ∼55 ppm and Cβ*_i–1_* of ∼62 ppm and Cβ of ∼69 ppm, typical of threonine residues for both the pre- [Figure S9] and post-fusion states [Figure S10]. Given the sequence of the LASV FD, this left one residue, L^266^, which is the only leucine residue to be preceded by a threonine residue, T^265^, as the most likely candidate for this assignment.

Nonetheless, before we delved too far into assigning an unknown number of conformers, we aimed to determine if there was any merit behind our notion that the LASV FD was adopting multiple conformations at a residue-specific level. Since we were confident that L^266^ had four distinct populations and several other leucine residues are distributed throughout the LASV FD, we decided to selectively ^15^N-label all leucine residues to further probe the existence of other multiple conformers and their population. A ^1^H – ^15^N HSQC of the isotopically labeled ^15^N–Leu FD at physiological pH [Figure 3A] and lysosomal pH with acidic bicelles [Figure 3B] correlated well with that of the entirely labeled FD. When we overlayed the ^1^H *–* ^15^N-Leu HSQC with the ^1^H *–* ^15^N HSQC and then looked at the corresponding strips in the different backbone experiments, we were led to the same ^1^H – ^15^N strips that we had previously identified as L^266^. This affirmed our notion that the observed strips with the same shifts arose from the same L^266^ residue and not a different residue with similar chemical shifts. From the ^1^H *–* ^15^N HSQC, we deduced the percentage of each population and assigned accordingly by taking the integration of each conformer. In the pre-fusion state, the first (a), second (b), third (c), and fourth (d) most populated conformer of L^266^ existed at 38%, 25%, 22%, and 15%, correspondingly [Figure 3C], which was comparable to the post-fusion state at 40%, 24%, 20%, and 16%, respectively [Figure 3D]. Notably, the PRE values and dynamics of L^266^ were somewhat analogous between each conformation, aside from the d conformer, indicative that each conformer was similarly exposed to the environment and flexible [Figure S11]. The difference observed here could be due to the d conformation being infrequently adopted and, thus, on a timescale that cannot be adequately captured by these experiments. Furthermore, we effectively applied this approach to the other leucine residues, i.e., L^280^, L^285^, and L^290^, allowing us to identify conformations for other leucine residues. We found two conformations for L^285^ and three for L^290^ in the pre-fusion state, but no additional conformers for the other leucine residues in the post-fusion state. For L^285^, the a conformation was occupied 68% of the time, whereas the b conformer was 32%. The a conformer of L^290^ was populated 53% of the time, b conformer was 34%, and c conformer was 13%. These findings indicate that multiple residues within the LASV FD adopt different populations, which can be different between the pre- and post-fusion states.

**Figure 3.**
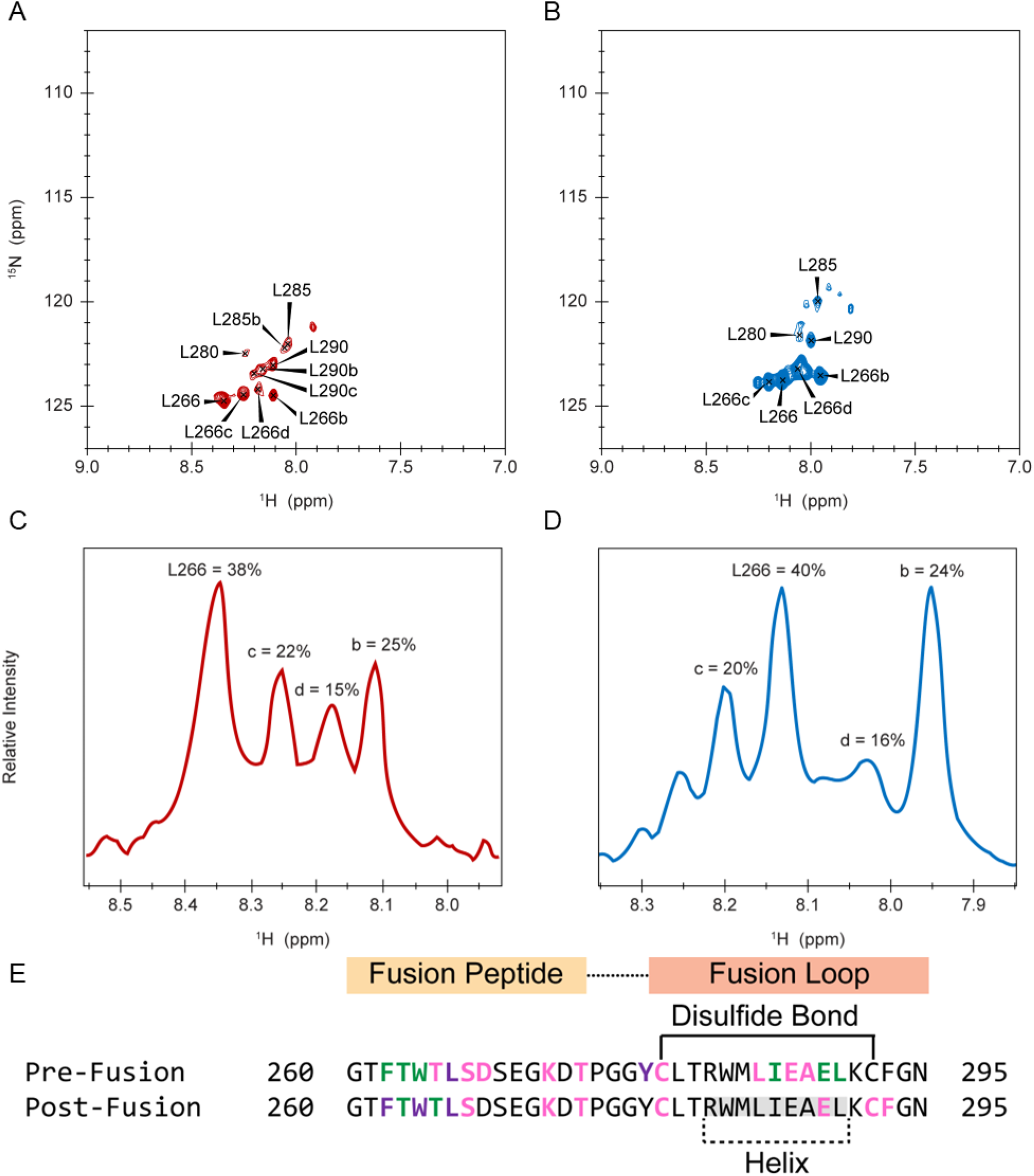
Multiple conformations are adopted by L^266^ in the LASV FD. [A,B] ^1^H - ^15^N-Leu HSQC spectra indicate a multitude of conformers exist for leucine residues in the [A] pre- and [B] post-fusion state. [C,D] Each L^266^ conformer is occupied for different percentages of time in the [C] pre- and [D] post-fusion state. All measurements were carried out on ∼1,000 µM of the LASV FD in 300 µL of 25 mM Na_2_HPO_4_, 100 mM NaCl, pH 7.0 at 20 °C (pre-fusion) or pH 4.0 with acidic bicelles, q = 0.5 at 45 °C (post-fusion). The most populated conformer (a) is the residue label, whereas the second (b), third (c), and fourth (d) most populated conformers are indicated as such. [E] Numerous residues throughout the FP and FL sample multiple conformations in the pre-fusion state (top), but only the FP continuing to do so in the post­fusion state (bottom). Residues with two conformers (pink), three conformers (green), and four conformers (purple) and the helix (grey box) formed in the post-fusion state are shown.

We next turned our attention back to the entire ^15^N-labelled FD to further elucidate residues with multiple populations [Table S6]. In the pre-fusion state, we found eight other residues in the LASV FD with two different conformations (T^265^, S^267^, D^268^, K^272^, T^274^, C^279^, E^287^, and A^288^), five with three conformations (F^262^, T^263^, W^264^, I^286^, and E^289^), and one with four conformations (Y^278^). Thus, a total of 17 residues (F^262^, T^263^, W^264^, T^265^, L^266^, S^267^, D^268^, K^272^, T^274^, Y^278^, C^279^, L^285^, I^286^, E^287^, A^288^, E^289^ and L^290^) with multiple conformations were identified in the pre-fusion state, equating to approximately 50% of the LASV FD having numerous populations [Figure 3E, top]. The FP exchanged populations slightly more than the FL with 9 out of 15 residues (60%) shown to have different conformers, as opposed to the 7 out of 17 residues (41%) in the FL. In the post-fusion state, however, the fraction of the LASV FD having different conformations decreased to 12 residues (33%) with a bulk majority of the same residues in the FP, but not the FL, continuing to exchange [Figure 3E, bottom]. Here, only four residues (24%) of the FL had two different conformers in the post-fusion state, including C^279^ and E^289^, which occurred in the pre-fusion state as well, and newly identified C^292^ and F^293^. Nonetheless, all residues identified in the FP to have multiple conformers in the pre-fusion state, except D^268^, continued to do the same in the post-fusion state. Intriguingly, while S^267^, K^272^, and T^274^ maintained two conformations, T^263^ and T^265^ now had three conformations, whereas F^262^ and W^264^ joined L^266^ in having four conformations. All of the residues identified to have multiple conformations occupied each population to a varying degree in the pre- and post-fusion state [Table S6]. On average, the a, b, c, and d conformers were populated 67 ± 19%, 24 ± 12%, 16 ± 6%, and 11 ± 6% of the time, accordingly, in the pre-fusion state. This remained relatively the same in the post-fusion state with the a, b, c, and d conformations adopted 67 ± 17%, 25 ± 11%, 16 ± 8%, and 10 ± 6% of the time, correspondingly.

Notably, the predicted dihedral angles of each conformer are relatively similar to each other in both the pre- [Table S7] and post-fusion [Table S8] states, which led us to question where these alternative conformers arose from. Thus, we employed site specific, ^19^F labeling of the LASV FD with 4-fluoro-D,L-phenylalanine, which has a fluorine attached at the para position of the aromatic. The rationale here is that fluorine is highly sensitive to its chemical environment, which allows us to monitor the side chain for chemical shifts undergone that occur in response to its surroundings^85^. Notably, there are two phenylalanine residues in the LASV FD: one at the N-terminal of the FP (F^262^) and one at the C-terminal of the FL (F^293^) [Figure 1]. Since F^262^, but not F^293^, appeared to sample numerous conformers in both the pre- and post-fusion state, we decided to selectively ^19^F-label F^262^. To accomplish this, we introduced a single mutation into F^293^ (i.e., F^293^W), such that only F^262^ would be labeled and analyzed. Given our previous data, we expected to see three peaks in the pre-fusion state and four peaks in the post-fusion state – accounting for one peak per conformer observed. Interestingly, our ^19^F experiments revealed that the side chain of F^262^ experienced two additional conformers in the pre-fusion state [Figure S12A] for a total of six conformers and one additional conformer in the post-fusion state [Figure S12B] for a total of five conformers. This suggests that these alternative conformers are due to the side chains of a given residue sampling different states. Altogether, it is evident that numerous residues throughout the LASV FD populate different conformers in the pre-fusion state, whereas only the FP continues to sample multiple conformations in the post-fusion state, likely because it is still solvent-exposed, but the FL becomes affixed when it associates with the membrane.

### A net negative charge akin to the lysosomal compartment correlates with increased LASV FD-initiated fusion, which is not influenced by positively charged residues

The LASV FD has been established to preferentially initiate fusion with the lysosomal membrane, which has a high propensity of anionic lipids that can impact fusion – as observed with other class one fusion viruses^43, 86–91^. We first aimed to determine if the LASV FD has a specificity for anionic lipids by performing a proof-of-concept titration with various molar ratios of POPC:POPG and, thus, various net negative charges, through the usage of isothermal titration calorimetry (ITC). We observed that the LASV FD had a weak affinity for vesicles that were completely zwitterionic and neutral (100 POPC) with a dissociation constant (K_d_) of 657 ± 17 μM [Figure 4A]. Minimal affinity was noted for vesicles with low concentrations of anionic lipids and negative charge (85:15 POPC:POPG) as denoted by the slight decrease in K_d_ to 542 ± 39 μM [Figure 4B]. In contrast, there was a drastic decrease in the K_d_ to 98.5 ± 19 μM when vesicles with an anionic lipid ratio and net negative charge approaching that of the lysosomal membrane were introduced (75:25 POPC:POPG), correlating with an increased affinity for the vesicles [Figure 4C]. This increase in affinity for the vesicles continued as the concentration of anionic lipids and, thus, net negative charge was increased to match that of the lysosomal compartment with a K_d_ of 11.7 ± 0.8 μM previously established for vesicles comprised of 65:35 POPC:POPG^41^. More specifically, a nearly 50-fold increase in affinity was witnessed between 85:15 POPC:POPG and 65:35 POPC:POPG vesicles, whereas an 8-fold increase was noted when the latter was compared to 75:25 POPC:POPG vesicles, indicating that anionic lipids influence LASV FD-initiated fusion.

**Figure 4.**
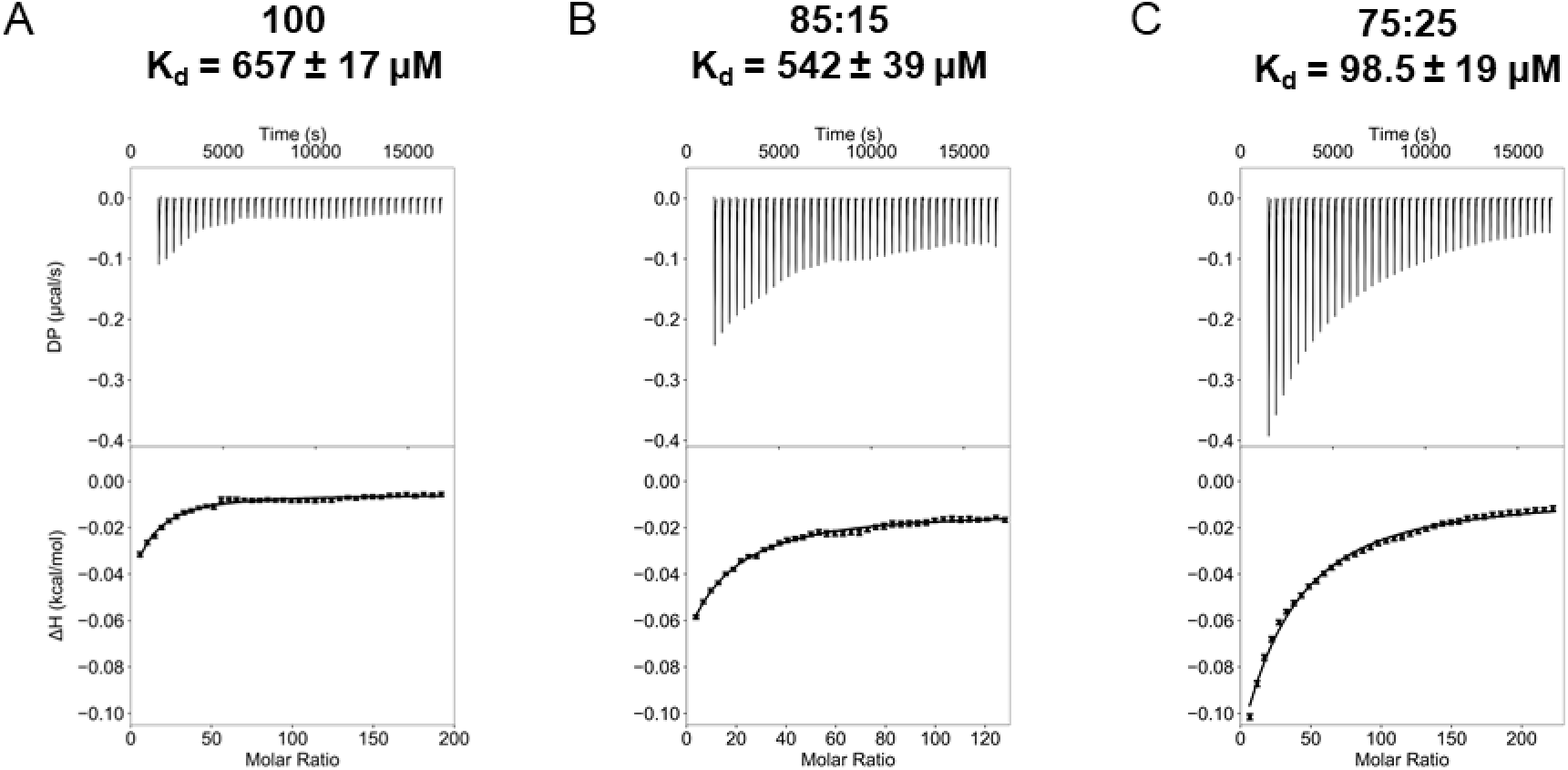
A net negative charge akin to the lysosomal membrane is important for the function of the LASV FD. [A] 100 POPC; [B] 85:15 POPC:POPG; and [C] 75:25 POPC:POPG were titrated into the protein. Displayed ratios are in terms of POPC:POPG. All ITC experiments were conducted in 10 mM NaOAc, 100 mM NaCI, pH 4.0 with vesicles titrated into the protein. Dissociation constants (K_d_) are displayed above the respective isotherm. An exemplary isotherm for LASV FD into 65:35 POPC:POPG is not shown as it has been published previously.

To investigate the structural and functional impact of the different POPC:POPG vesicles on the LASV FD, we employed CD spectroscopy and a FRET-based lipid mixing assay, respectively. We found that the helical propensity of the FD was dependent upon the concentration of POPG present [Figure 5A]. The LASV FD appeared to exist in a predominantly random coil conformation in vesicles comprised of 100 POPC and 85:15 POPC:POPG, as delineated by the single dip at 200 nm. As the concentration of anionic lipids and, thus, net negative charge, was increased to 75:25 POPC:POPG and 65:35 POPC:POPG, the secondary structure of the LASV FD transitioned to be mainly helical, indicated by the double dip at 208 and 222 nm, with similar helical content at 33% and 35%, correspondingly. This correlated with an exponential increase in fusion, as expected given that the helical conformation of the LASV FD has been established to be its fusogenic form [Figure 5B]. In particular, we witnessed no fusion in 100 POPC vesicles. Minimal fusion was noted in 85:15 POPC:POPG vesicles, which was followed by a negligible increase in 75:25 POPC:POPG vesicles. In contrast, fusion readily occurred in vesicles comprised of 65:35 POPC:POPG. Despite the FD having comparable helical propensity in 75:25 POPC:POPG and 65:35 POPC:POPG, a 5-fold increase in fusion was observed between 75:25 POPC:POPG and 65:35 POPC:POPG vesicles. These findings suggest that the LASV FD is sensitive to the concentration of anionic lipids present with optimal fusion occurring when a net negative charge akin to the lysosomal membrane is present.

**Figure 5.**
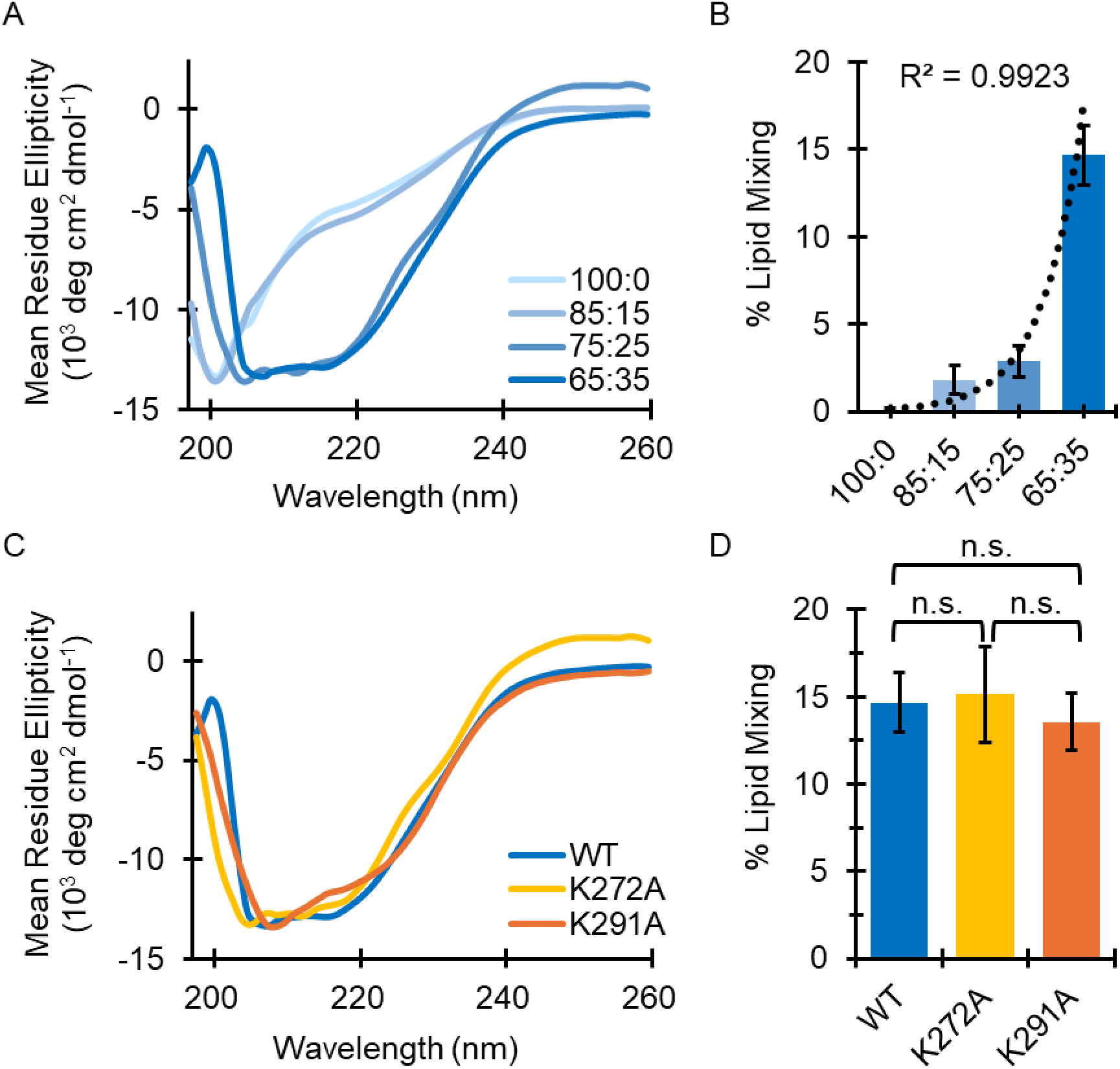
Structure and function of the LASV FD is reliant on the overall negative charge, which is not mediated by positively charged lysine residues. [A] Global secondary structure is dependent on the zwitterionic:anionic lipid ratio with a helical structure adopted only once a net negative charge akin to the lysosomal membrane was achieved. [B] Exponential relationship between increased concentrations of POPG and fusion with optimal fusion occurring in 65:35 POPC:POPG (n ≥ 16). [C] In vesicles comprised of 65:35 POPC:POPG, all lysine mutants (K272A, yellow; K291A, orange) had similar secondary structure to WT (blue). [D] Mutation of the lysine residues had no consequential impact on fusion (n ≥ 15). Student’s t-Test assuming unequal variances used to calculate the P-value; n.s. = not significant. Displayed ratios are in terms of POPC:POPG.

We next asked if this preference for a high propensity of negatively charged lipids was mediated by an ionic interaction. A sequence alignment revealed that there are three positively charged residues within the LASV FD: K^272^, located in the FP, and R^282^ and K^291^, found in the FL [Figure 1]. Notably, K^272^ is not conserved, R^282^ is moderately conserved, and K^291^ is completely conserved. Single point mutations were generated to alter the positively charged side chains of residues K^272^ and K^291^ into a chemically inert alanine (i.e., K^272^A and K^291^A) instead. Mutagenesis of R^282^ was not performed as it has previously been established to be a part of a salt bridge that is important to LASV FD-initiated fusion^41^. Surprisingly, the secondary structure of the LASV FD remained relatively unchanged when either residue was mutated, as defined by the deep dips at 208 and 222 nm [Figure 5C]. To be more specific, the helical content of K^272^A and K^291^A was 32% and 33%, respectively, which was akin to that observed in WT (35%)^41^. There were no significant impacts on function with both K^272^A and K^291^A having similar amounts of fusion as WT [Figure 5D]. This corresponded to a high affinity for the membrane with a K_d_ of 19.8 ± 1.0 μM for K^272^A [Figure S13A] and 19.2 ± 1.2 μM for K^291^A [Figure S13B]. Thus, the observed preference for a negatively charged lipid does not appear to be mediated by an ionic interaction between the membrane and the LASV FD.

### The LASV FD has an enhanced interaction with BMP, an anionic and conical lipid prevalent within the lysosomal membrane

We next sought to deduce if anionic lipids with physiological relevance to eukaryotic cells, chiefly those found in the lysosomal compartment, have a particular effect on the LASV FD. The lysosomal membrane has a high composition of lipids known to influence fusion, namely bis(monoacylglycero)phosphate (BMP), phosphatidylethanolamine (PE), and phosphatidylserine (PS). We thus conducted experiments with 65:35 POPC:X where X was either BMP, POPE, or POPS. Notably, BMP has both a negatively charged lipid head group, similar to POPS, and an inverted cone shape, akin to POPE, which is not captured when studying POPS and POPE alone. As such, we also investigated the effect that vesicles comprised of 65:17.5:17.5 POPC:POPS:POPE had on the function of the LASV FD. We demonstrate that the FD only has helical content when in the presence of POPC:BMP, which was comparable to that observed in POPC:POPG [Figure 6A]. More specifically, even at 2X higher concentrations of POPE (i.e., 65:35 POPC:POPE vs 20% PE in the lysosomal compartment) or 10X higher concentrations of POPS (i.e., 65:35 POPC:POPS vs 2% PS in the lysosomal compartment) than physiologically relevant, the LASV FD seemed to exist as a random coil structure, as the presence of a deep minimum near 200 nm was witnessed. This remained true in the presence of POPS and POPE together (POPC:POPS:POPE), as the FD still existed in a largely random coil conformation. However, in BMP, dips at 208 and 222 nm were observed, indicative of a helix. While the 208 nm intensity was comparable to that observed in POPC:POPG, there was a slight decrease in the 222 nm intensity, correlating with a decrease in helical content to 30% and potentially one to two fewer residues involved in the helical structure.

**Figure 6.**
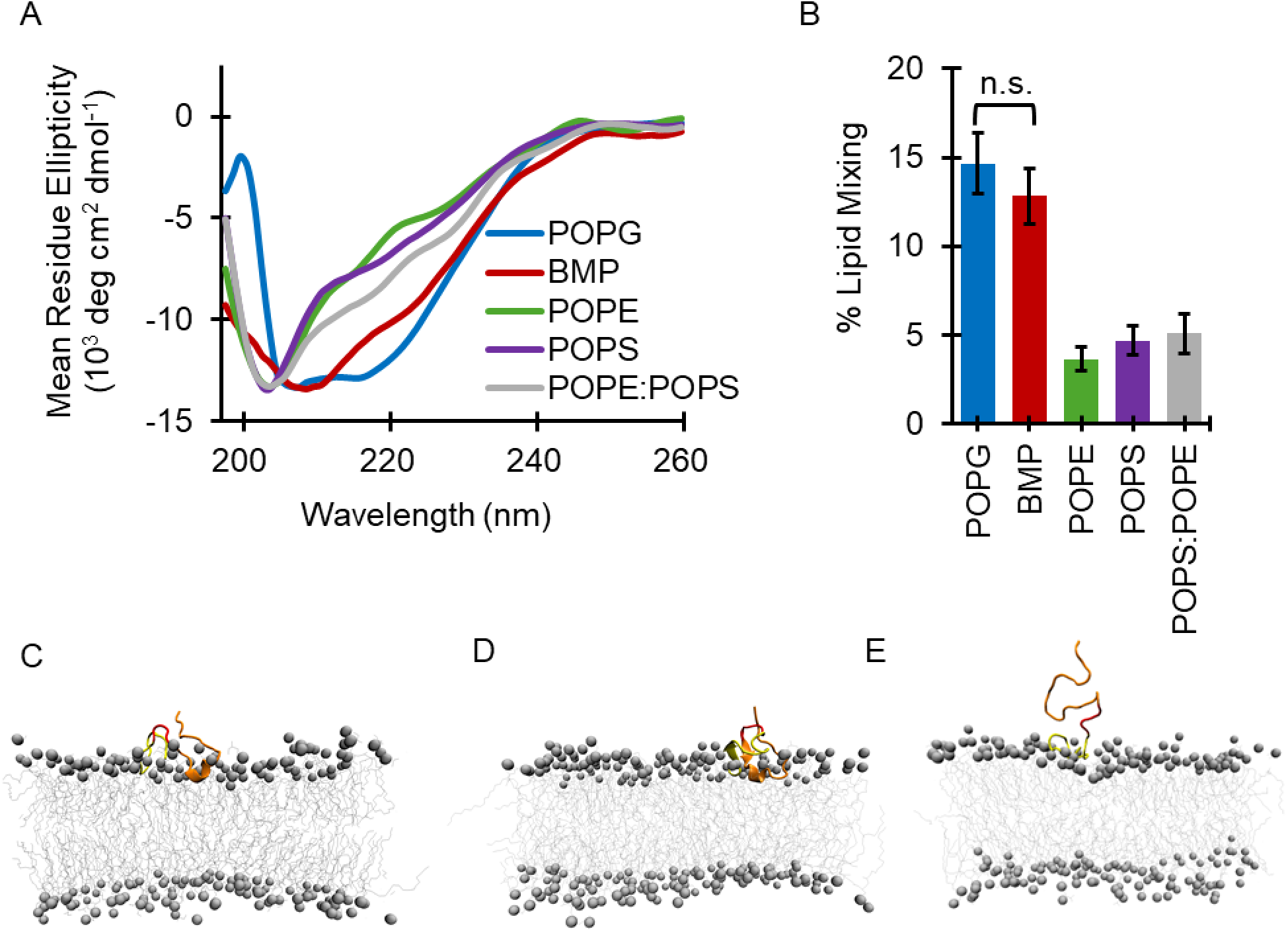
Anionic lipids, chiefly BMP, impact the structure and function of the LASV FD. [A] Global secondary structure was influenced by the type of physiologically relevant lipid present with coil character observed in all lipids (POPE, green; POPS, purple; POPS:POPE, grey), except BMP (red), which appeared similarly to POPG (blue). [B] Fusion of the LASV FD was dependent on the lipid present with optimal fusion occurring in BMP (n ≥ 12). Student’s t-Test assuming unequal variances used to calculate the P-value. All results were significant unless designated; n.s. = not significant. [C-E] After 2 µs, the LASV FD adopts a helix in its FL (orange) that associates with the lipid head group, whereas the FP (yellow) remains solvent exposed in membranes comprised of [C] POPC:POPG and [D] POPC:BMP, but not [E] POPC:POPS.

Since there is a direct link between the structure and initiation of fusion, we aimed to decipher how these physiologically relevant lipids impacted the function of the LASV FD. We witnessed a negligible decrease in binding affinity and fusion when the FD was in POPC:BMP as opposed to POPC:POPG [Figure 6B, S14A]. The FD had a K_d_ of 25.2 ± 3.7 μM in vesicles comprised of BMP, which was 2-fold higher than that established in POPC:POPG; however, there was no statistically significant difference in fusion. In contrast, when the LASV FD was in vesicles comprised of POPC:POPE, POPC:POPS, or POPC:POPS:POPE, we witnessed a nearly identical reduction in binding affinity and fusion when compared to POPC:BMP and POPC:POPG. The FD had a minimal affinity for vesicles comprised of POPC:POPE, as denoted by the drastic increase in the K_d_ to 606 ± 155 μM [Figure S14B], which led to an over 3-fold decrease in fusion. This remained unchanged in vesicles containing POPC:POPS or POPC:POPS:POPE as the K_d_ was 453 ± 32 μM [Figure S14C] and 415 ± 21 μM [Figure S14D], respectively, with similar amounts of fusion. It could be argued that studying POPS and POPE together did not truly encapsulate the double unsaturation and anionic head group at concentrations similar to BMP, which is why vesicles comprised of 65:17.5:17.5 POPC:POPS:POPE did not have similar levels of LASV FD-initiated fusion witnessed in 65:35 POPC:BMP. However, we saw no significant difference in the ability of the LASV FD to initiate fusion in lipids with the same head group, but different tail saturations [Figure S15]. Thus, the functionality of the FD has a clear preference on the type of lipid present, namely BMP.

Molecular dynamics (MD) simulations were employed to visualize the interaction of the FD with various membranes, specifically the degree to which the helical structure existed and, thus, penetrated the membrane. Three membrane systems were simulated, including POPC:POPG, POPC:BMP, and POPC:POPS. Systems with POPC:POPE and POPC:POPS:POPE were not simulated since the FD has a clear preference for anionic lipids and there was no significant difference experimentally in the structure or function when compared to POPS. We provide evidence that the LASV FD could associate with membranes comprised of POPC:POPG [Figure 6C; Video S1] and POPC:BMP [Figure 6D; Video S2]. Helical content was observed in POPC:POPG [Figure S16A] and POPC:BMP [Figure S16B], especially in the FL for residues R^282^ – L^290^, but more residues were frequently involved in the formation of the helix in POPC:POPG than POPC:BMP, supporting our experimental CD data. The helix inserted itself just below the lipid head group in both the POPC:POPG and POPC:BMP systems. On the contrary, the FD did not associate whatsoever with the POPC:POPS membrane [Figure 6E; Video S3] and remained largely random coil over the entire course of the simulation [Figure S16C]. Put together, our results indicate that anionic lipids in the lysosomal membrane, chiefly BMP, are imperative for the LASV FD to interact with the membrane and initiate fusion.

## Discussion

Infection with LASV results in Lassa fever – a severe hemorrhagic fever with no FDA-approved therapeutic options. Delivery of LASV’s genetic material is achieved by the FD, which undergoes a conformational change to become helical and initiate fusion at the lysosomal compartment. In other class one fusion proteins, the FD is either an FP or FL, but in LASV, the FD has both an FP and FL in tandem [Figure 1]. Previous work has demonstrated that the LASV FD preferentially initiates fusion at the lysosomal membrane, at which point it adopts a fusogenic, helical conformation^41^. However, the structural and functional characteristics governing this interaction remain poorly understood, especially the location of the helix and the role of particular lipids in the lysosomal membrane. Here, we reveal that the FL of the LASV FD forms a helix to associate with the membrane, whereas the FP remains solvent-exposed and samples multiple conformations. We indicate that the LASV FD has a preference to initiate fusion with anionic lipids, namely BMP, which is not mediated by an ionic interaction with lysine residues located in the FP or FL. These observations highlight the distinct roles of the FP and FL in facilitating membrane fusion and suggest a unique mechanism for viral infectivity.

In the literature, the FD of class one fusion proteins undergo conformational changes, often becoming helical, to penetrate the host cell membrane and mediate fusion. For example, the HIV FD adopts an amphipathic helix upon membrane insertion, contributing to the destabilization of the target host cell membrane such that fusion can occur^29–31, 92–94^. The influenza FD must also undertake a similar structure transition to mediate fusion, but instead adopts a helical hairpin structure^32–36, 95, 96^. While the LASV FD undergoes a pH-dependent conformational change from the pre- [Figure 2A; Table S2] to post-fusion [Figure 2B; Table S3] state, our results suggest that the FP remains solvent-exposed while the FL has helical content and is the primary membrane-interacting component [Figure 2D]. More specifically, a helix is adopted for residues R^282^ – L^290^ within the FL [Figure 2C], which is juxtaposed to a properly formed disulfide bond between C^279^ and C^290^ [Figure S4, S5] that is important for the tertiary structure and initial interaction of the FD with the host cell membrane^41^. We speculate that this disulfide bond serves to stabilize the helix in the membrane and increase the energy required to remove it^44, 49, 97^. This agrees with the EBOV FD that mediates fusion through a hydrophobic fist that is clamped by a disulfide bond^28, 37, 38, 42, 98, 99^. However, while the EBOV FD inserts itself deeply within the membrane, our PRE experiments indicate that the LASV FD associates with the membrane more superficially, just beneath the lipid head groups [Figure 2D, S7; Table S4]. Our dynamics experiments agreed as the LASV FD became more restricted once fusion occurred, especially the FL [Figure 2E, S8; Table S5]. This aligns more closely with the SARS-CoV-2 FD, which also has both an FP and FL in tandem. In the SARS-CoV-2 FD, the FP adopts a boomerang structure, akin to the influenza FD, and becomes more restricted after association below the lipid head groups. In contrast, the FL remains exposed to the environment and dynamic, serving as a mechanical stabilizer for the FP and interacting with extracellular factors to increase fusogenicity^45–48^. Thus, we suggest that the FP and FL of the LASV FD have opposite roles as their counterparts in the SARS-CoV-2 FD. The FL forms a boat-like structure that is capped by a disulfide bond and facilitates the initial interaction with the membrane, while the FP, especially residues D^268^ – T^274^, functions as its mechanical stabilizer and is like a sail for the boat [Figure 7].

**Figure 7.**
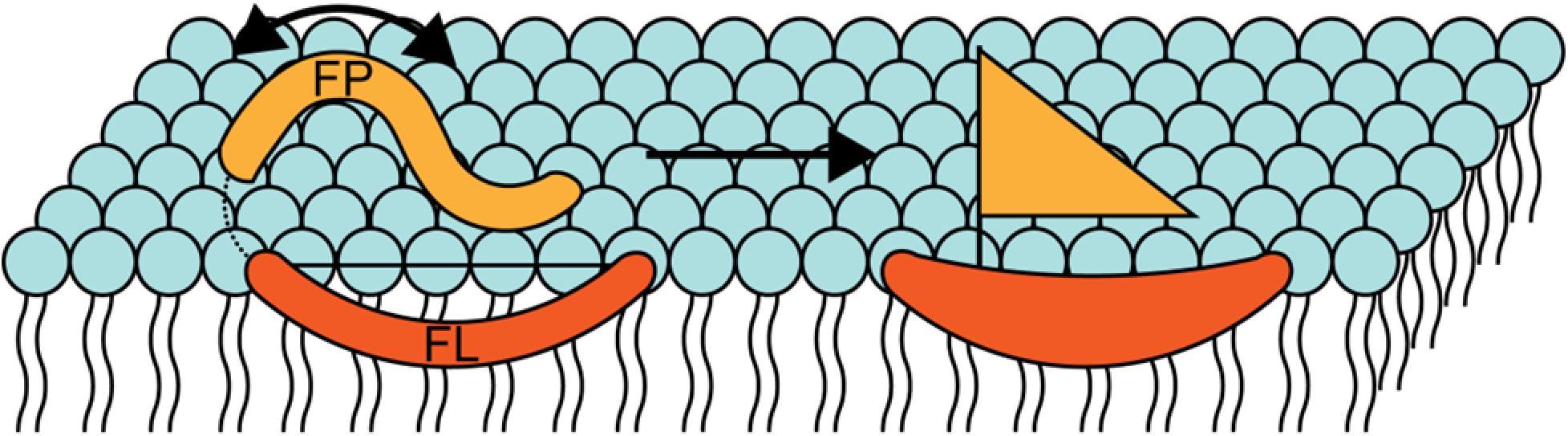
Structural model of the LASV FD associated with the host cell membrane. The Fl (orange, C^279^- N^295^) adopts a helix (R^282^ - L^290^) that is capped by a disulfide bond (black line, C^279^ and C^292^) and associates with the membrane in a shallow manner, almost like a boat. The FP (yellow, G^260^ - T^274^) remains solvent exposed, especially the residues (D^268^ - T^274^) closest to the linker region (dotted line, P^275^ - Y^278^), and adopts multiple conformations (double-ended arrow) to engage with the additional extracellular factors, acting as a sail for the boat.

During our structural investigation, we uncovered multiple peaks with different ^1^H – ^15^N shifts, but identical Cα, Cβ, and CO shifts in both the pre- [Figure S9] and post-fusion [Figure S10] states. One plausible explanation would be that a single residue adopted multiple conformations on a slow enough timescale, i.e., milliseconds to seconds, that could be captured via solution NMR spectroscopy. For example, recent studies have revealed alternative conformations undergone by the influenza FD on a sub-millisecond scale and suggested structural plasticity^32, 33, 100–109^. A three-model system was thus developed to explain these distinct conformers as each had a different tertiary structure that uniquely influenced fusion pore formation. This aligns with a new theory that has recently emerged as structural intermediates have been described for various class one and two fusion proteins, which are believed to properly orient their FDs for membrane remodeling^110–113^. We postulate that the LASV FD also follows a multisystem model to carry out fusion with up to four different conformations [Figure 3A,B]; although, a much wider array of structural transitions may be quickly sampled, and, thus, not captured during our NMR experiments. This likely affects the side chains more than the backbone as the predicted dihedral angles were relatively similar between the different conformers in both the pre-[Table S7] and post-fusion [Table S8] states. Our ^19^F experiments, which are particularly sensitive to environmental changes, support this notion as additional conformers were observed in both the pre- [Figure S12A] and post-fusion [Figure S12B] states when the side chain of F^262^ was specifically labeled at the para position of the aromatic. In the pre-fusion state, roughly half of the LASV FD transiently samples multiple conformations with residues distributed throughout both the FP (9 residues, 60%) and FL (7 residues, 40%) having different conformers [Figure 3C; Table S6]. We theorize that this large sampling of alternative conformations by the LASV FD in the pre-fusion state explains the small, but uniform, reduction in the overall signal intensity upon Gd-DTPA titration [Figure S6]. A similar phenomenon has been observed in intrinsically disordered proteins (IDPs) where the Gd-DTPA probe does not form a complex with the IDP but rather becomes homogeneously distributed around the IDP^114-116^. IDPs are notoriously extremely flexible, often functioning without a well-defined structure until folding, which is often binding-induced. In the pre-fusion state, we contend that the LASV FD behaves similarly to an IDP and slowly samples alternative conformations to extensively interact with various environmental factors. Once the right conditions are sensed, the FD is triggered to transition into the post-fusion state so fusion occurs at the appropriate location.

Interestingly, in the post-fusion state, the FP (8 residues, 53%) continued to effectively adopt alternative conformations, whereas the FL (4 residues, 24%) was largely limited to a single conformer [Figure 3D; Table 4.5]. We theorize that the FP continued to sample multiple conformations and remained dynamic in the post-fusion state as it was still solvent-exposed [Figure 2D, S7; Table S4]. This is in alignment with our notion that adopting multiple conformations likely allows for more interactions with the environment to increase fusion. In contrast, the FL undergoes binding-induced folding and is “locked” into a single conformer, which provides a rationalization for why only the peripheral residues continued to transiently populate alternate conformers in the post-fusion state. The formation of the helix within the FL thus potentially functions as the rate-limiting step for the multisystem model of the LASV FD, akin to the influenza FD. It is important to note that one of these residues within the FL, E^289^, is part of a salt bridge shown to significantly influence the structure and function of the LASV FD^41^. In influenza, it has been postulated that the conformational change between the different structures is driven by changes in the protonation states of the acidic side chains at low pH and is an early rate-limiting step^109, 117–120^. We believe that protonation of E^289^ results in a similar occurrence and helps to drive the transition of the FL into its single, post-fusion conformer. This is in agreement with literature where the protonation of E^289^ at a low pH results in the formation of a new bonding network that prompts the adoption of the fusogenic, helical conformation^41^. Notably, this includes the bonding network with R^282^, the salt bridge counterpart to E^289^, which is also important for the structure and function of the LASV FD, but to a lesser extent than E^289^. Taken together, this indicates that the LASV FD follows a multisystem model to mediate fusion, potentially exploiting its conformational plasticity to have optimal fusion under varying physiological conditions [Figure 7].

One of the most important, but often overlooked, environmental factors in membrane fusion is the lipid composition of the target host cell membrane. A key feature of LASV FD-initiated fusion is its preference for the lysosomal compartment, where the fusogenic, helical conformation is only adopted at a low pH of 4.0^24–27, 41, 121, 122^. The predominant lipids of the lysosomal membrane are phosphatidylcholine (PC), phosphatidylethanolamine (PE), bis(monoacylglycero)phosphate (BMP), sphingomyelin (SM), and cholesterol (CHOL) at a molar ratio of 50:20:15:10:5, respectively^90^. Additionally, there are trace amounts of the anionic lipids phosphatidylinositol (PI) and phosphatidylserine (PS) at roughly 2-3% and 1-2%, respectively, but totaling no more than 5%. BMP is an anionic lipid that is specific to the lysosome and is the most abundant anionic lipid present in this organelle. Moreover, PE is unusual in that it does not carry a charge but has an inverted cone shape similar to that argued to exist within BMP, which is known to induce negative curvature in lipid membranes and elicit fusion more readily^86–88, 91^. Therefore, this led us to theorize that the LASV FD interacts with specific lipids, namely anionic lipids, to influence fusion. We show that increased concentrations of POPG, a rudimentary anionic lipid, resulted in a progressively higher affinity of the FD for the lipid bilayer [Figure 4A-C]. More specifically, the FD existed in its nonfusogenic, random coil form in vesicles containing little to no concentrations of POPG, but transitioned into its fusogenic, helical form as the concentration of POPG was increased [Figure 5A], which correlated to a positive, exponential increase in fusion [Figure 5B]. We postulate that while a low pH may be necessary for fusion to occur, it in itself is insufficient to trigger a robust interaction between the LASV FD and the target host cell membrane. Collectively, this gives credence to our hypothesis that anionic lipids impart a specific function within LASV FD-initiated fusion.

A preference for anionic lipids has been linked to positively charged residues in several class one fusion proteins, including HIV and SARS-CoV-2^43, 50, 123^. Given the similarities between the LASV FD and SARS-CoV-2 FD, we decided to investigate if positively charged residues in the LASV FD contributed to the anionic lipid preference. A sequence alignment of the LASV FD alongside other arenaviruses revealed three positively charged residues: K^272^, R^282^, and K^291^ [Figure 1]. K^272^ is located in the FP and not conserved, whereas R^282^ and K^291^ were both located in the FL and conserved to different degrees. Single mutants were created such that K^272^ and K^291^ were converted to a chemically inert alanine (i.e., K^272^A and K^291^A), while R^282^ was not mutated as its necessity for the structure and function of the LASV FD was previously established. We hypothesized that K^291^ would be most likely to have an ionic interaction with the anionic lipids given its position in the lipid head group, as demonstrated via our NMR experiments [Figure 2D, S7], and degree of conservation. In turn, we believed that a lack of charge at this position would impede the necessary ionic interaction and result in decreased function and/or structural perturbations. On the other hand, mutation of K^272^ would have no significant impact on the function or structure of the LASV FD as it remains solvent-exposed throughout the fusion process. Unexpectedly, we observed no significant structural change in either mutant when compared to WT [Figure 5C]. Subsequently, the binding affinity [Figure S13A,B] and fusogenicity [Figure 5D] of both mutants were similar to WT. These findings thus suggest that the LASV FD does not rely on ionic interactions to mediate its interaction with anionic lipids but rather exploits lipid specific properties to facilitate fusion.

Anionic lipids, particularly BMP, have been implicated to have a positive and specific influence on fusion for several other viruses that employ the endocytic pathway, such as influenza, SARS-CoV-2, and vesicular stomatitis virus^43, 87–89, 124^. For example, studies on the entry of flaviviruses, which include class two fusion proteins like dengue virus and Zika virus, indicate that their FDs preferentially interact with anionic lipids, including BMP, to promote viral fusion^125–127^. We hypothesized that a similar preference might influence LASV FD-initiated fusion, especially since the lysosomal compartment has a high composition of BMP and the FD is specifically influenced by the property of a lipid itself. We also postulated that POPS and POPE could impart a particular interaction as well given that they are also known to influence fusion. Surprisingly, we found that the FD had a significantly higher affinity for BMP over the other anionic lipids [Figure S14A-D]. Additionally, helical content [Figure 6A] and, thus, fusion [Figure 6B] were only noted in BMP and similar to that seen in POPG. MD simulations reinforced these observations, demonstrating that the LASV FD remains helical and membrane-associated in a lipid environment with POPG [Figure 6C, S16A; Video S1] and BMP [Figure 6D, S16B; Video S2], but becomes more disordered in the presence of POPS [Figure 6E, S16C; Video S3]. It could be claimed that this observation was due to a difference in the tail saturation between POPS and BMP and the head group charge between POPE and BMP. However, both of these properties should have been captured when POPS and POPE were in tandem, yet we observed no significant difference in the structure or function of the LASV FD when POPS and POPE were in combination or alone. Even when we introduced vesicles with the same lipid head groups of PG, PS, or phosphatidic acid (PA), but different tail saturations (i.e., single unsaturation (POXX, 16:0 – 18:1) vs double unsaturation (DOXX, 18:1)), we saw no significant difference in fusion [Figure S15]. A previous study on the complete LASV GPC also indicated a specific effect on fusion mediated by BMP and not POPS or DOPS^86^. Based on our findings, a possible explanation is that LASV FD-initiated fusion has a specificity for a particular type of anionic head group moiety. In the literature, it has been shown that the head group of BMP is the only mammalian glycerophospholipid to have a distinctive sn-1;sn-1’ arrangement, as opposed to the traditional sn-3:sn-1’ orientation, that confers increased stability and is found in PG^128, 129^. This unique configuration has been found to directly influence fusion by the SARS-CoV-2 FD and likely explains why we observed some fusion in POPA/DOPA, which has partial similarity to BMP and PG^43^. Nonetheless, the preference of the SARS-CoV-2 FD for BMP is a result of both the unique bonding network for the head group moiety and decreased lipid packing caused by the increased unsaturation of the tails, which is not true for the LASV FD^49^. Put together, these findings enforce the idea that LASV FD-initiated fusion is optimized for the unique composition of the lysosomal membrane, with a distinct preference for the unique head group moiety found in BMP.

In summary, our study provides new insights into the structural plasticity of the LASV FD and its role in viral entry. We have illustrated that conformational changes occur from the pre- to post-fusion state with the FP and FL having distinct structural and function roles. Here, the ability of the FD to sample multiple conformations may be a key evolutionary advantage that allows it to fine-tune its fusion process in response to different lipid environments and initiate fusion at the appropriate time. Additionally, the low pH of the lysosomal compartment alone is necessary, but insufficient to trigger fusion with the target host cell membrane and requires anionic lipids. Positively charged residues within the FD do not impart a specific, ionic interaction with anionic lipids. Instead, the LASV FD takes advantage of the membrane disorder created by BMP to preferentially interact with the membrane and initiate fusion. Overall, we have provided a deeper understanding of the molecular details governing the structure and function of the LASV FD. These findings highlight the critical role of both the conformational flexibility of the FD and anionic lipids, chiefly BMP, in fusion and suggest that disrupting either of these interactions could serve as a novel antiviral strategy. Further research is warranted to explore the implications of these alternative conformations and the role of lipids in a broader context of related arenaviruses, potentially providing insights into viral evolution and therapeutic targets.

## Accession Codes

The associated NMR chemical shifts have been deposited in the BMRB under the accession numbers 53085 and 53084 for the pre- and post-fusion states, correspondingly.

## Author Contributions

H.N.P and J.L. designed the experiments. H.N.P. performed the experiments and was supported by K.A. and Q.M.M. H.N.P. analyzed the data. S.S. and W.I. performed and analyzed the molecular dynamics simulations. H.N.P and J.L. prepared the manuscript.

## Notes

The authors declare no conflict of interest.

## Acknowledgments

We would like to thank all lab members for their assistance in editing this manuscript.

