## Supplemental Document for "The Lassa Virus Fusion Domain has structural plasticity and exploits bis(monoacylglycero)phosphate for fusion"

| Page | Supporting Information |
|---|---|
| S1 | Structures of purchased lipids |
| S2 | Temperature titration of the LASV FD in the pre- and post-fusion states via solution NMR spectroscopy |
| S3 | Chemical shift indexing of LASV FD vs BMRB values |
| S4 | Cβ strips for the pre- and post-fusion states of the LASV FD |
| S5 | Standard curve for free thiol present in LASV FD under native and reducing conditions |
| S6 | Gd-DTPA titration of the LASV FD in the pre-fusion state |
| S7 | Gd-DTPA and 16-DSA titration of the LASV FD in the post-fusion state |
| S8 | $R_1$ and $R_2$ relaxation rates of the LASV FD in the pre- and post-fusion states |
| S9 | $^1H$ – $^{15}N$ strips for $L^{266}$ in the pre-fusion state |
| S10 | $^1H$ – $^{15}N$ strips for $L^{266}$ in the post-fusion state |
| S11 | Membrane depth and dynamics of different $L^{266}$ conformations |
| S12 | $^{19}F$ spectra of $F^{293}W$ in the pre- and post-fusion state |
| S13 | Exemplary isotherms of the lysine mutants |
| S14 | Exemplary isotherms of the LASV FD in different lipids |
| S15 | Fusion of the LASV FD in lipids with different tail saturations |
| S16 | Helical graphs of the LASV FD in POPG, BMP, and POPS |

| | |
|---|---|
| S17 | Lipid compositions of the membrane systems generated for MD simulations |
| S18 | TALOS+ statistics for the pre-fusion state |
| S19 | TALOS+ statistics for the post-fusion state |
| S21 | Average relative intensities of LASV FD in different paramagnetic probes |
| S22 | Overall relaxation times of the LASV FD in the pre- and post-fusion state |
| S23 | Populations for residues with multiple conformers in the pre- and post-fusion state |
| S24 | Predicted dihedral angles for different conformers in the pre-fusion state |
| S25 | Predicted dihedral angles for different conformers in the post-fusion state |

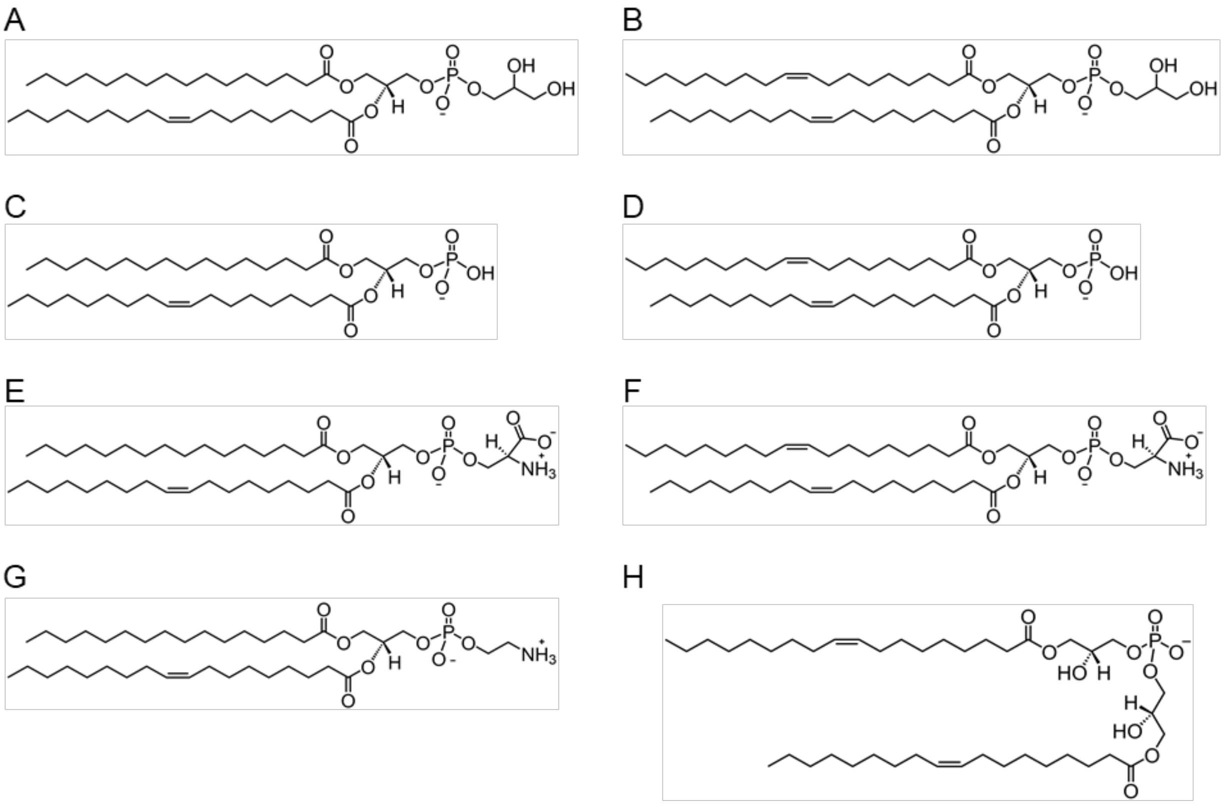

**Figure S1.** Structure of various lipids employed in this study that were purchased from Avanti Lipids. [A] 16:0–18:1 1-palmitoyl-2-oleoyl-*sn*-glycero-3-phospho-[1′-rac-glycerol] (POPG); [B] 1,2-dioleoyl-*sn*-glycero-3-[phospho-rac-(3-lysyl(1-glycerol))] (DOPG); [C] 1-palmitoyl-2-oleoyl-*sn*-glycero-3-phosphate (POPA); [D] 1,2-dioleoyl-*sn*-glycero-3-phosphate (DOPA); [E] 1-palmitoyl-2-oleoyl-*sn*-glycero-3-phospho-L-serine (POPS); [F] 1,2-dioleoyl-*sn*-glycero-3-phospho-L-serine (DOPS); [G] 1-palmitoyl-2-oleoyl-*sn*-glycero-3-phosphoethanolamine (POPE); and [H] bis(monooleoylglycero)phosphate (S,R Isomer) (BMP). All structures drawn in ChemDraw.

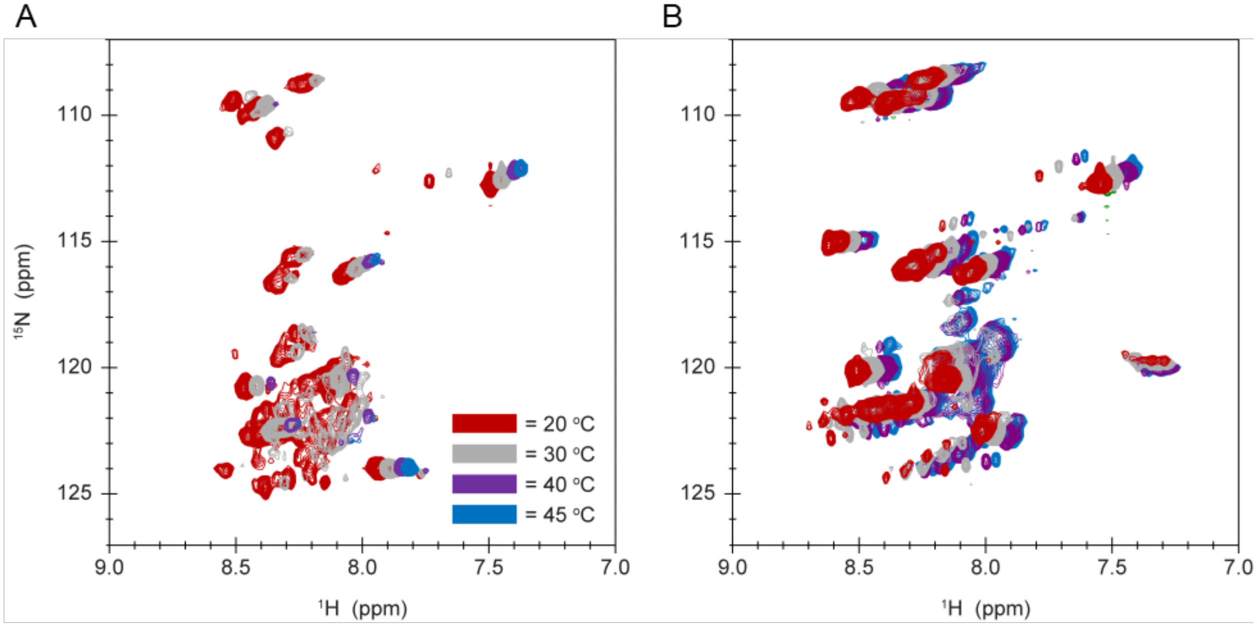


**Figure S2.** The LASV FD requires a different temperature for the different fusion states to be effectively visualized via NMR spectroscopy. [A,B] An overlay of 1H – 15N HSQC spectra for the [A] pre- and [B] post-fusion state of the LASV FD at 20 ℃ (red), 30 ℃ (grey), 40 ℃ (purple), and 45 ℃ (blue). All measurements were carried out on ~500 μM of the LASV FD in 300 μL of 25 mM Na2HPO4, 100 mM NaCl, pH 7.0 (pre-fusion) or pH 4.0 with acidic bicelles, q = 0.5 (post-fusion).

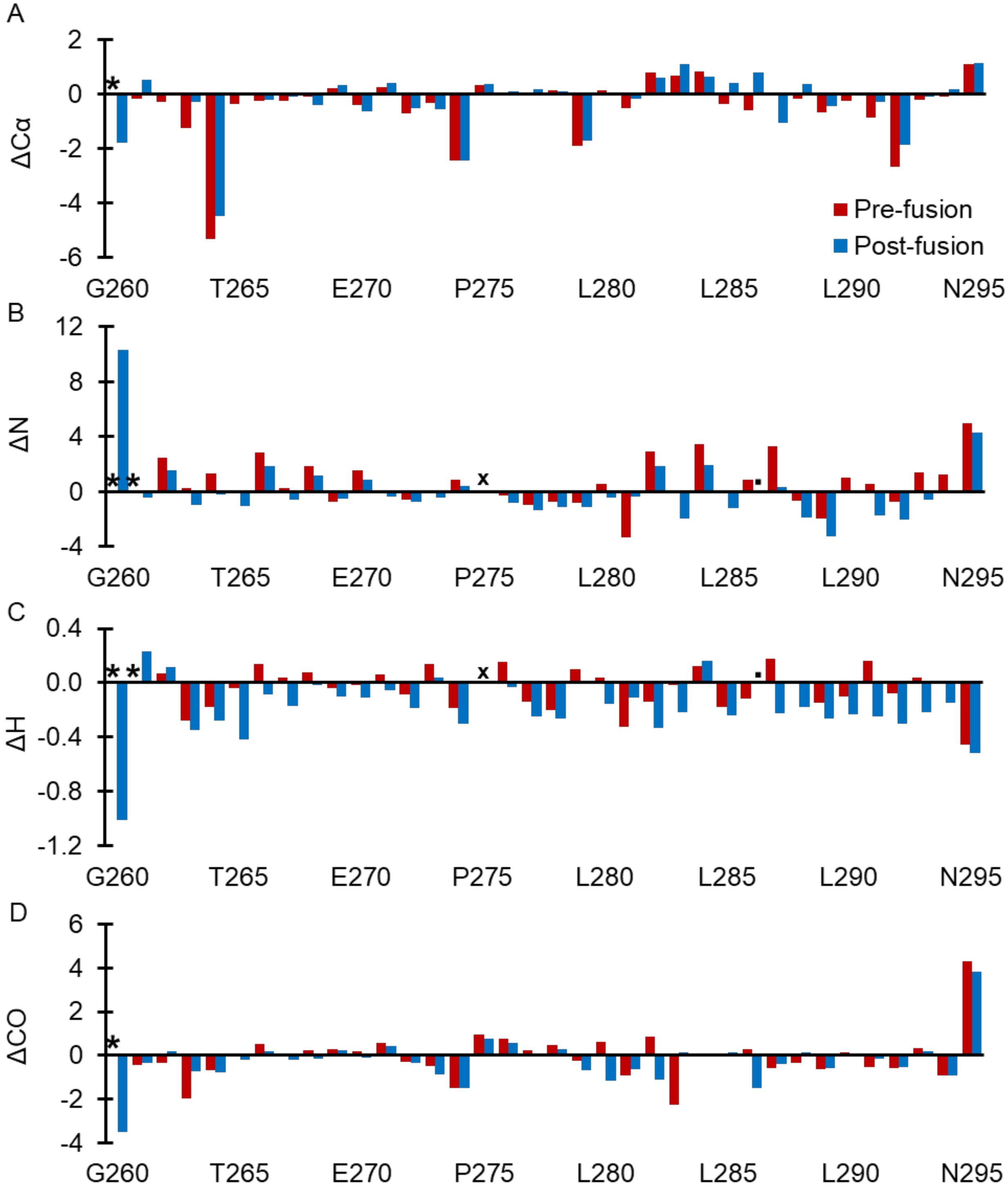


**Figure S3.** Chemical shift indexing in comparison to backbone atoms does not reveal information regarding the secondary structure of the LASV FD. [A] ΔCα [B] ΔN; [C] ΔH; and [D] ΔCO. All measurements were carried out on ~500 μM of the LASV FD in 300 μL of 25 mM Na2HPO4, 100 mM NaCl, pH 7.0 at 20 ℃ (pre-fusion) or pH 4.0 with acidic bicelles, q = 0.5 at 45 ℃ (post-fusion). Proline residues and other residues that could not be assigned for both states are marked (X), whereas residues that could not be assigned for the pre- or post-fusion state are marked with an asterisk (*) or period (.), accordingly.

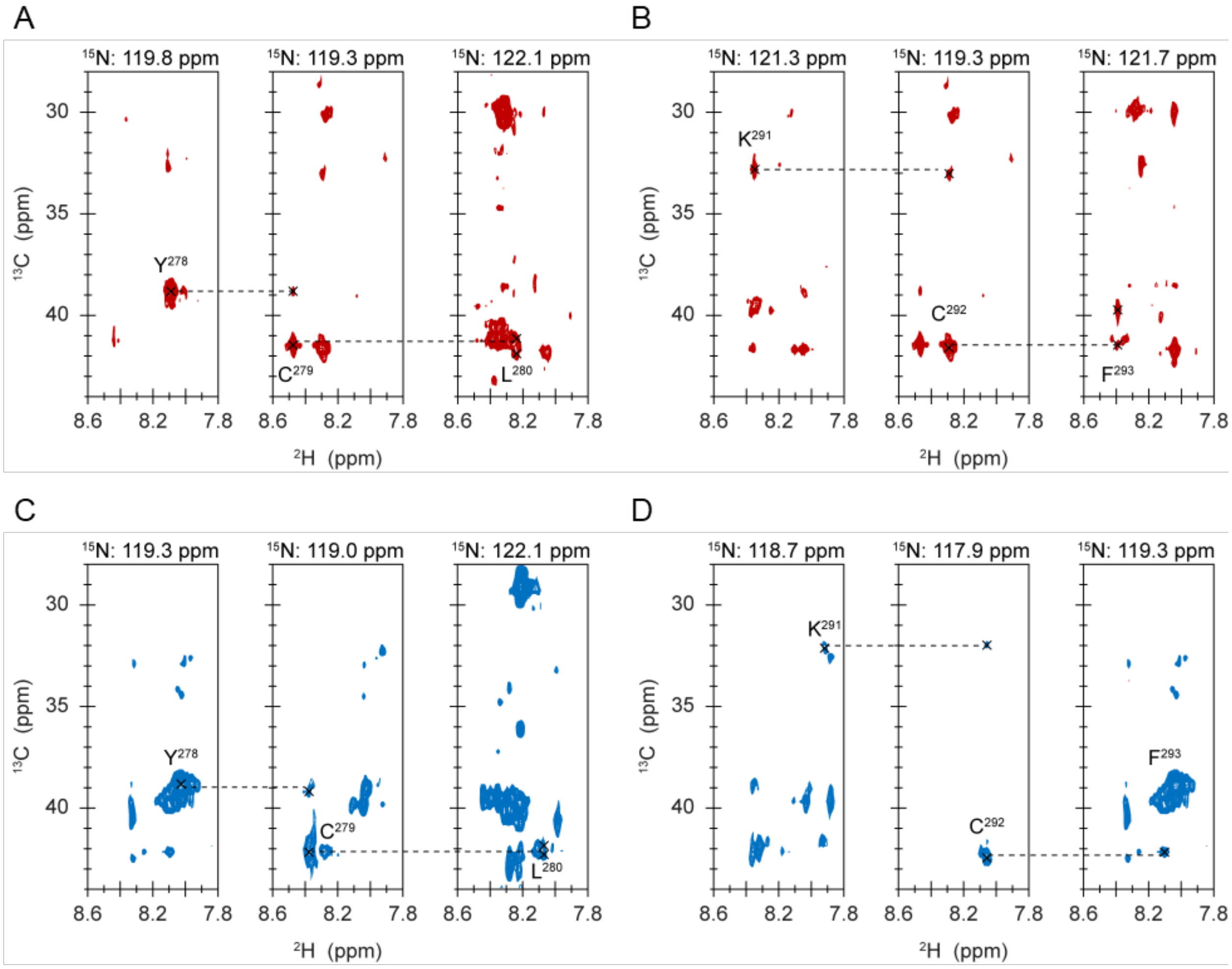


**Figure S4.** Cβ shifts confirm the presence of an internal disulfide bond within the FL of the LASV FD in both the pre- and post-fusion states. [A] Pre-fusion C279; [B] Pre-fusion C292; [C] Post-fusion C279; and [D] Post-fusion C292 all have 13Cβ chemical shifts greater than 35 ppm, indicative that the cysteine residues were oxidized. All data were acquired from HN(CA)CB experiments with 650 μM of a triple labelled sample (2H/13C/15N) in 25 mM Na2HPO4, 100 mM NaCl, pH 7.0 at 20 ℃ [A, B] or pH 4.0 at 45 ℃ with acidic bicelles, q = 0.5 [C,D].

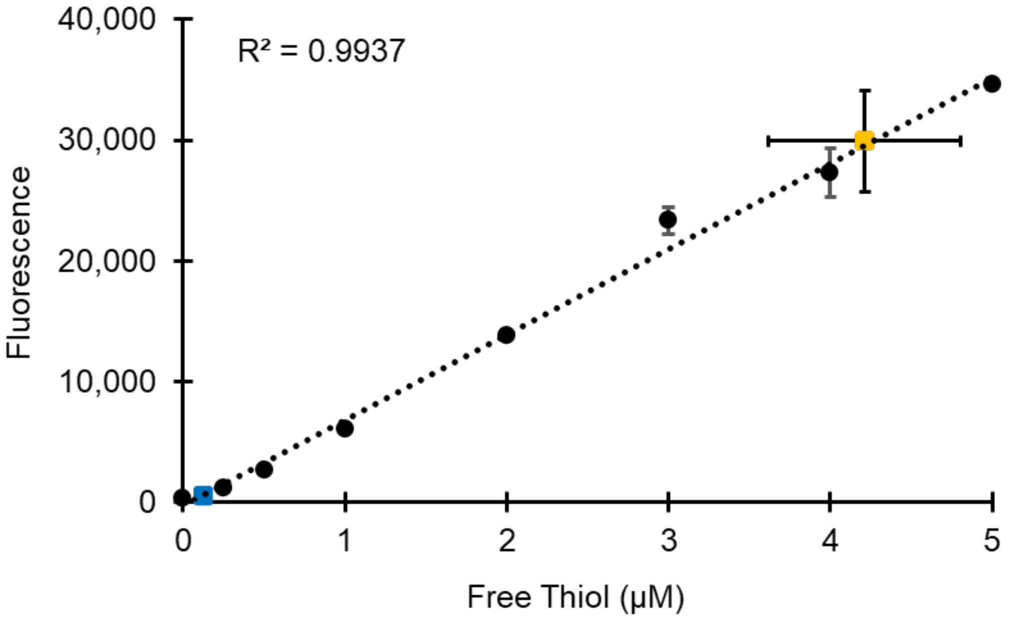


**Figure S5.** A large majority of the LASV FD has a properly formed disulfide bond. Under native conditions (blue, square; n = 3), the LASV FD had low concentrations of free thiol, suggestive that the disulfide bond was properly formed. Upon introduction of 1 mM TCEP to the FD (yellow, square; n = 2), there was a large increase in the concentration of free thiol present, indicative that the disulfide bond was disrupted. Assay was performed according to the manufacturer's protocol with standards in duplicates. Standard curve (black, circle) was generated to determine the unknown thiol concentration. The average fluorescence of 1 mM TCEP was run and subtracted from the average fluorescence of the FD with 1 mM TCEP. Errors were propagated accordingly from the standard deviations.

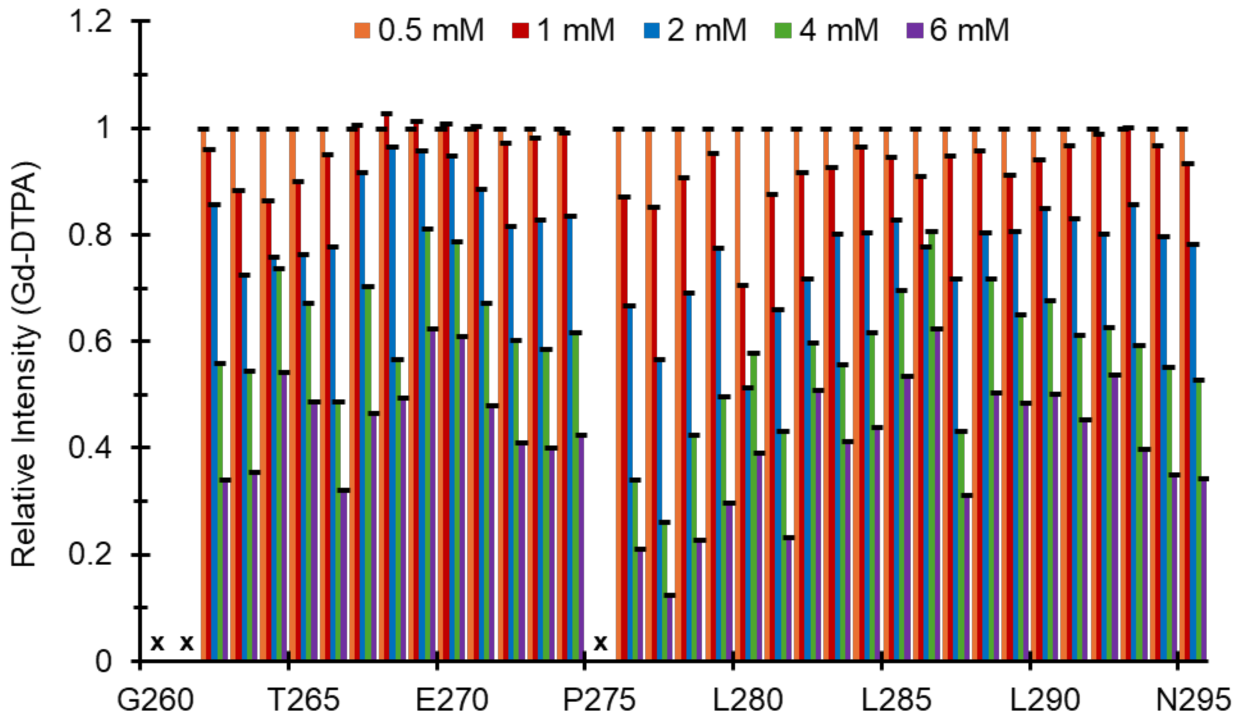


**Figure S6.** The LASV FD is solvent-exposed in the pre-fusion state. Progressive titration of the water-soluble, paramagnetic agent Gd-DTPA in increments of 0.5 mM (orange), 1 mM (red), 2 mM (blue), 4 mM (green), and 6 mM (blue) into the LASV FD revealed that all residues experience virtually the same amount of quenching. The relative intensity prior to titration is not shown due to increasing signals from the paramagnetic effect. Data normalized to 0.5 mM Gd-DTPA for each titration. Error bars shown are propagated from the signal-to-noise ratio and standard error of the mean (SEM). All measurements were carried out on ~500 µM of the LASV FD in 300 µL of 25 mM Na2HPO4, 100 mM NaCl, pH 7.0 at 20 ℃. Proline residues and other residues that could not be assigned are marked (X).

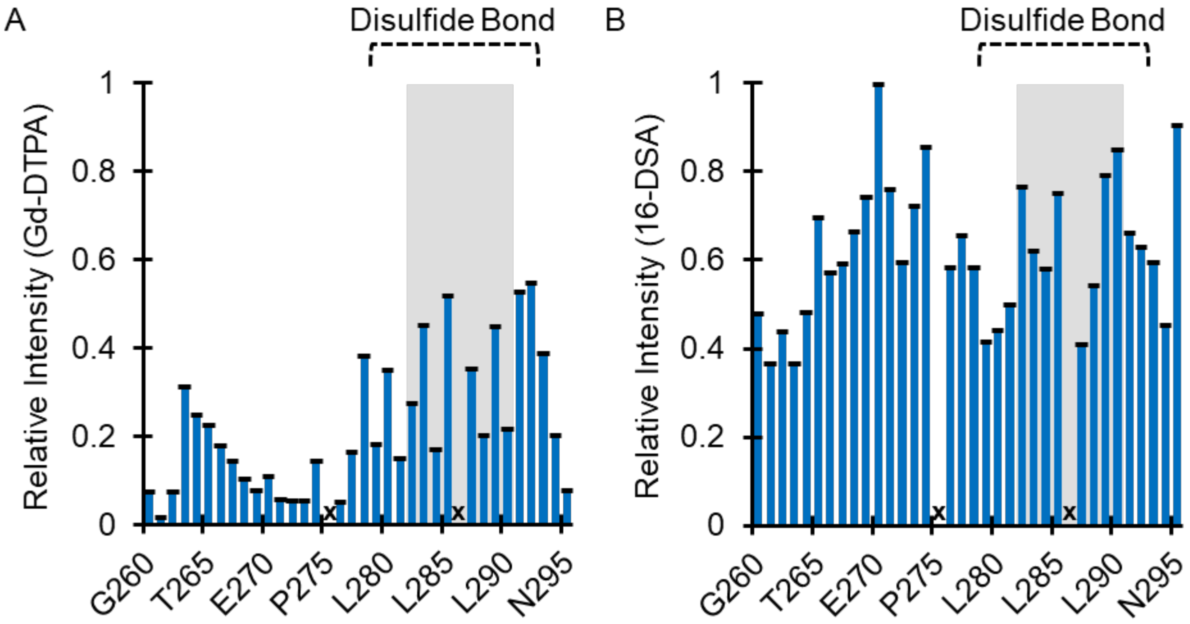


**Figure S7.** In the post-fusion state, the LASV FD associates with the membrane in a shallow manner via its FL while the FP remains solvent-exposed. [A] Introduction of 6 mM Gd-DTPA into the system containing the LASV FD resulted in significant quenching of the LASV FP, particularly D268 – T274, but not the FL. Titration of 6 mM 16-DSA into the LASV resulted in nearly identical quenching for both the FP and FL. For all experiments, the relative intensity at 0.5 mM of a given probe was subtracted due to increasing signals. All error bars shown are propagated from the signal-to-noise ratio and standard error of the mean (SEM). All measurements were carried out on ~500 µM of the LASV FD in 300 µL of 25 mM Na2HPO4, 100 mM NaCl, pH 4.0 with acidic bicelles, q = 0.5 at 45 ℃. The disulfide bond ($C^{279}$ and $C^{292}$, dashed line) and helix ($R^{282}$ – $L^{290}$, transparent grey box) are indicated. Proline residues and other residues that could not be assigned are marked (X).

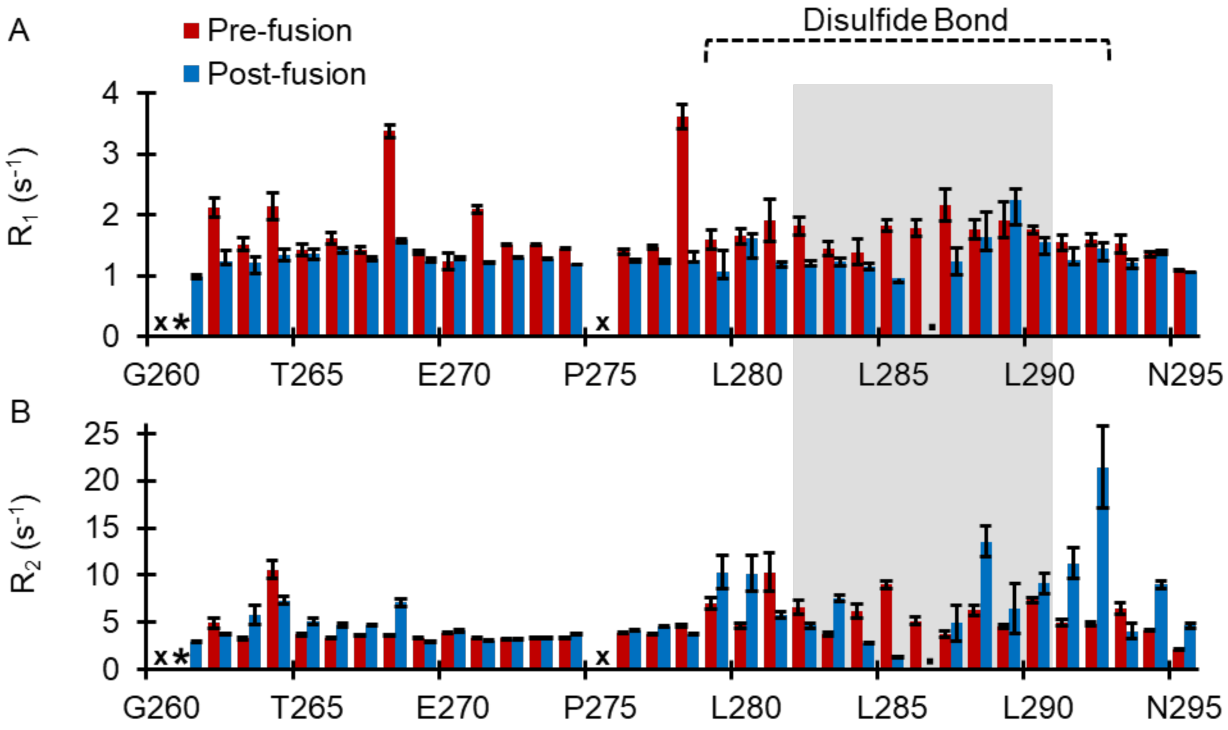


**Figure S8.** Dynamic properties were slightly altered in several regions within the LASV FD from the pre- (red) to post-fusion (blue) state. [A] Both the FP and FL have similar $R_1$ relaxation rates in the pre-fusion state that were similar but decreased in the post-fusion state. [B] $R_2$ relaxation rates for the FP were similar between the pre- and post-fusion states but increased for the FL in the post-fusion state. Error bars shown are propagated from the signal-to-noise ratio. All measurements were carried out on ~500 µM of the LASV FD in 300 µL of 25 mM $Na_2HPO_4$, 100 mM NaCl, pH 7.0 at 20 °C (pre-fusion) or pH 4.0 with acidic bicelles, q = 0.5 at 45 °C (post-fusion). The disulfide bond ($C^{279}$ and $C^{292}$, dashed line) and helix ($R^{282}$ – $L^{290}$, transparent grey box) are indicated. Proline residues and other residues that could not be assigned for both states are marked (X), whereas residues that could not be assigned for the pre- or post-fusion state are marked with an asterisk (*) or period (.), accordingly.

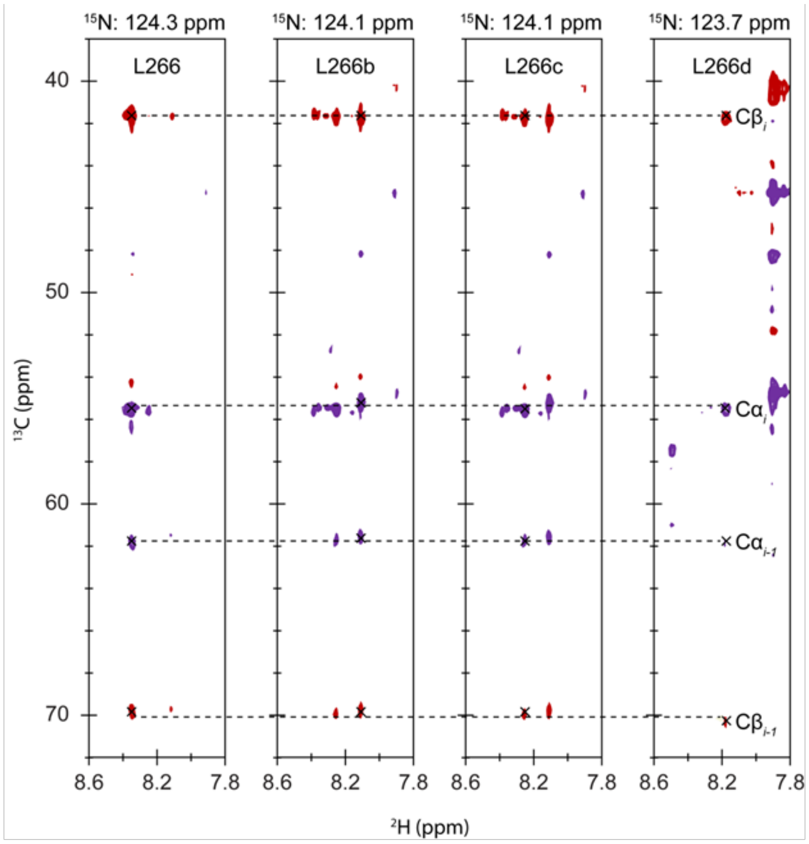


**Figure S9.** Multiple conformations for L[266] were observed in the pre-fusion states. Four different $^1H$ –$^{15}N$ strips had the same Cα (purple) and Cβ (red) shifts for the *i* and *i – 1*, corresponding to L[266] and T[265], accordingly. Data was acquired from HN(CA)CB experiments with 650 µM of a triple labelled sample ($^2H/^{13}C/^{15}N$) in 25 mM $Na_2HPO_4$, 100 mM NaCl, pH 7.0 at 20 ℃. The most populated conformer (a) is the residue label, whereas the second (b), third (c), and fourth (d) most populated conformers are indicated as such.

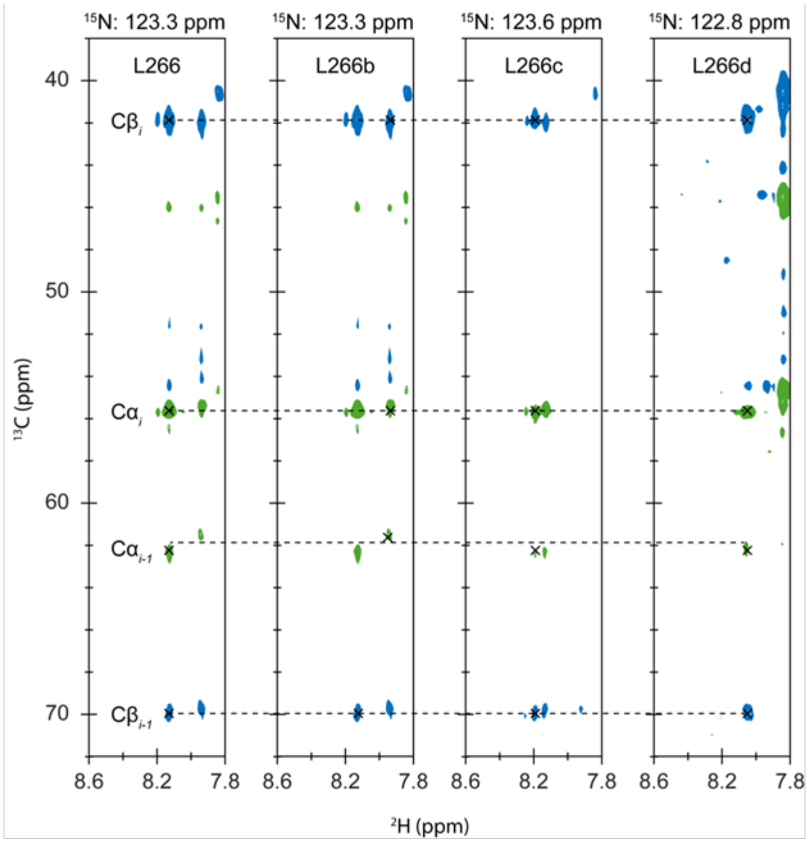


**Figure S10.** $L^{266}$ has several conformers in the post-fusion state. Different $^{15}N$ strips with the same Cα (green) and Cβ (blue) shifts were observed in the post-fusion state for $L^{266}$. All data were acquired from HN(CA)CB experiments with 650 μM of a triple labelled sample ($^{2}H/^{13}C/^{15}N$) in 25 mM $Na_2HPO_4$, 100 mM NaCl, pH 4.0 at 45 ℃ with acidic bicelles, q =0.5. The most populated conformer (a) is the residue label, whereas the second (b), third (c), and fourth (d) most populated conformers are indicated as such.

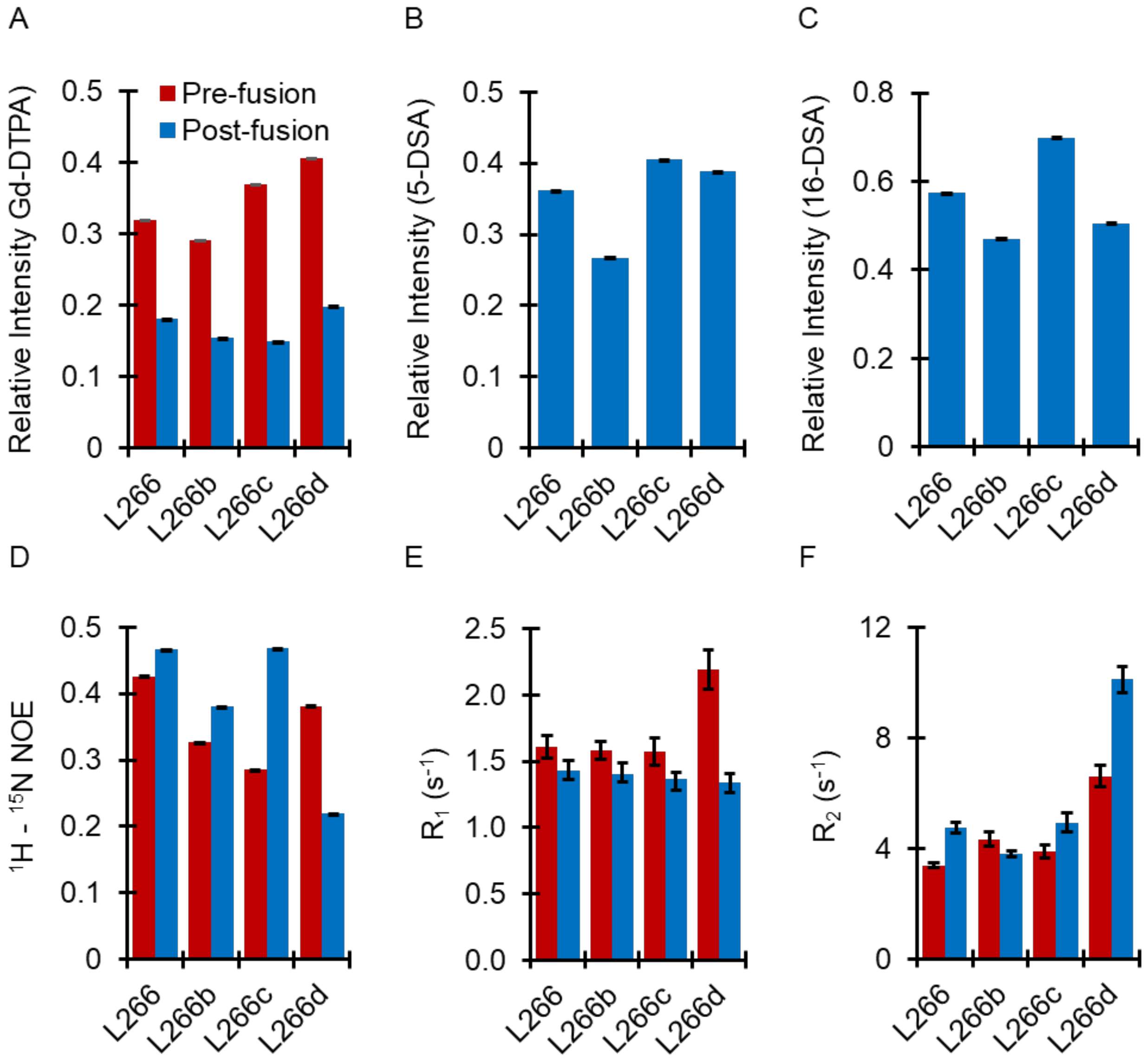


**Figure S11.** Membrane depth and dynamics of $L^{266}$ are relatively similar between the different conformations. [A-C] Each conformer of $L^{266}$ experiences comparable amounts of quenching in the presence of [A] 6 mM Gd-DTPA, [B] 8 mM 5-DSA, and [C] 6 mM 16-DSA. [D-E] The different $L^{266}$ populations have similar dynamics as revealed by [D] $^{1}H$ – $^{15}N$ NOE, [E] $R_1$ relaxation, and [F] $R_2$ relaxation experiments. Error bars shown are propagated from the signal-to-noise ratio. All measurements were carried out on ~500 μM of the LASV FD in 300 μL of 25 mM $Na_2HPO_4$, 100 mM NaCl, pH 7.0 at 20 ℃ (pre-fusion, red) or pH 4.0 with acidic bicelles, q = 0.5 at 45 ℃ (post-fusion, blue). The second (b), third (c), and fourth (d) most populated conformers of a given residue are indicated.

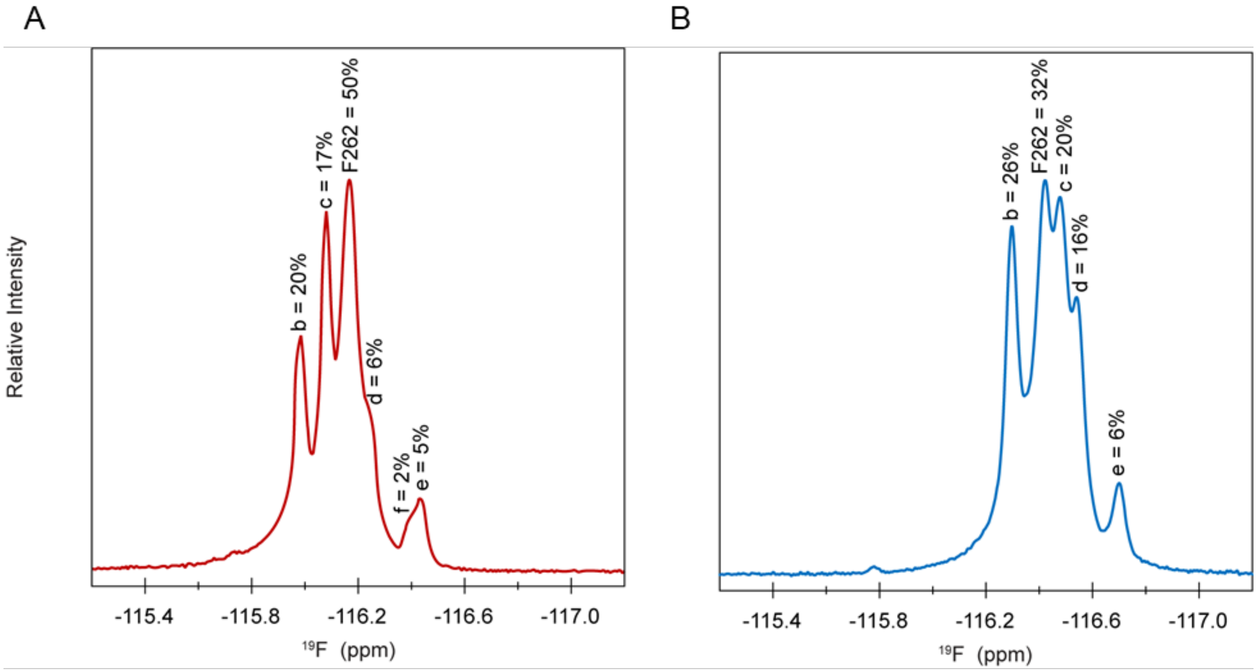


**Figure S12.** Site specific $^{19}F$ labeling of $F^{262}$ in the LASV FD revealed additional conformers adopted by the side chain that were not observed in the backbone assignment. [A,B] Each $F^{262}$ conformer is occupied for different percentages of time in the [A] pre- and [B] post-fusion state. All measurements were carried out on ~1,000 µM of $F^{293}W$ in 300 µL of 25 mM $Na_2HPO_4$, 100 mM NaCl, pH 7.0 at 20 ℃ (pre-fusion) or pH 4.0 with acidic bicelles, q = 0.5 at 45 ℃ (post-fusion). The most populated conformer (a) is the residue label, whereas the second (b), third (c), fourth (d), fifth (e), and sixth (f) most populated conformers are indicated as such.

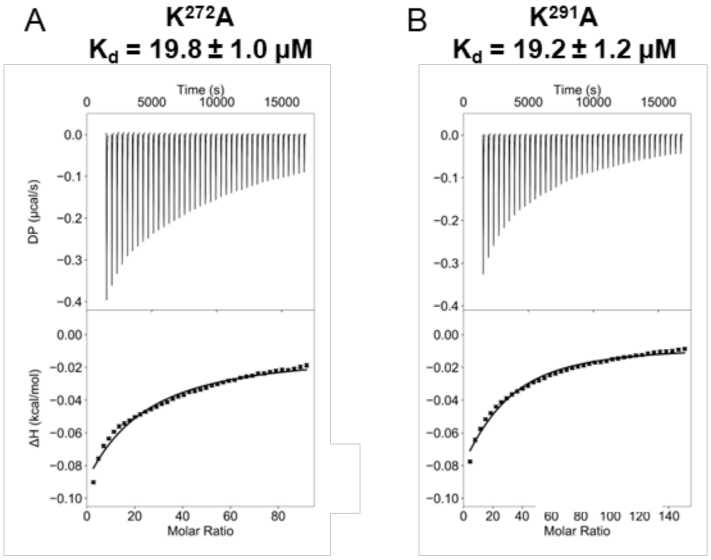


**Figure S13.** Mutation of the charged lysine residue located within the FP ($K^{272}$) or FL ($K^{291}$) to a chemically inert alanine residue had no impact on the binding affinity of the LASV FD. [A] $K^{272}$A and [B] $K^{291}$A mutant FD. Both ITC experiments were conducted in 10 mM NaOAc, 100 mM NaCl, pH 4.0 with 65:35 POPC:POPG vesicles titrated into the protein. Dissociation constants ($K_d$) are displayed above the respective isotherm.

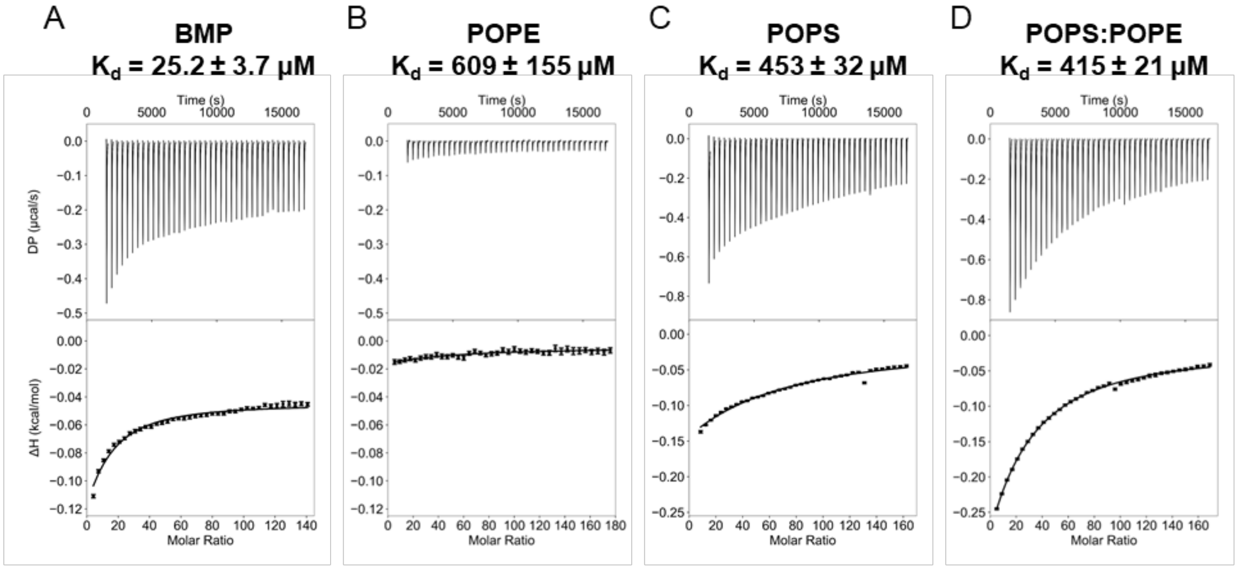


**Figure S14.** The LASV FD has the highest binding affinity for vesicles comprised of BMP. [A] 65:35 POPC:BMP; [B] 65:35 POPC:POPE; [C] 65:35 POPC:POPS; and [D] 65:17.5:17.5 POPC:POPS:POPE were titrated into the protein. All ITC experiments were conducted in 10 mM NaOAc, 100 mM NaCl, pH 4.0 with vesicles titrated into the protein. Dissociation constants ($K_d$) are displayed above the respective isotherm.

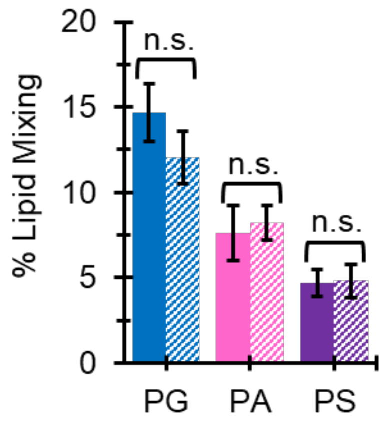


**Figure S15.** Tail saturation has no impact on LASV FD-initiated fusion. Lipids with a single unsaturated tail (POXX, solid color) and double unsaturated tail (DOXX, dashed color) did not have significant impact on the ability of the LASV FD to initiate fusion if the same head group moiety was present, i.e., PG (blue), PA (pink), and PS (purple) (n ≥ 9). Student's t–test assuming unequal variances used to calculate the *P*-value; n.s. = not significant.

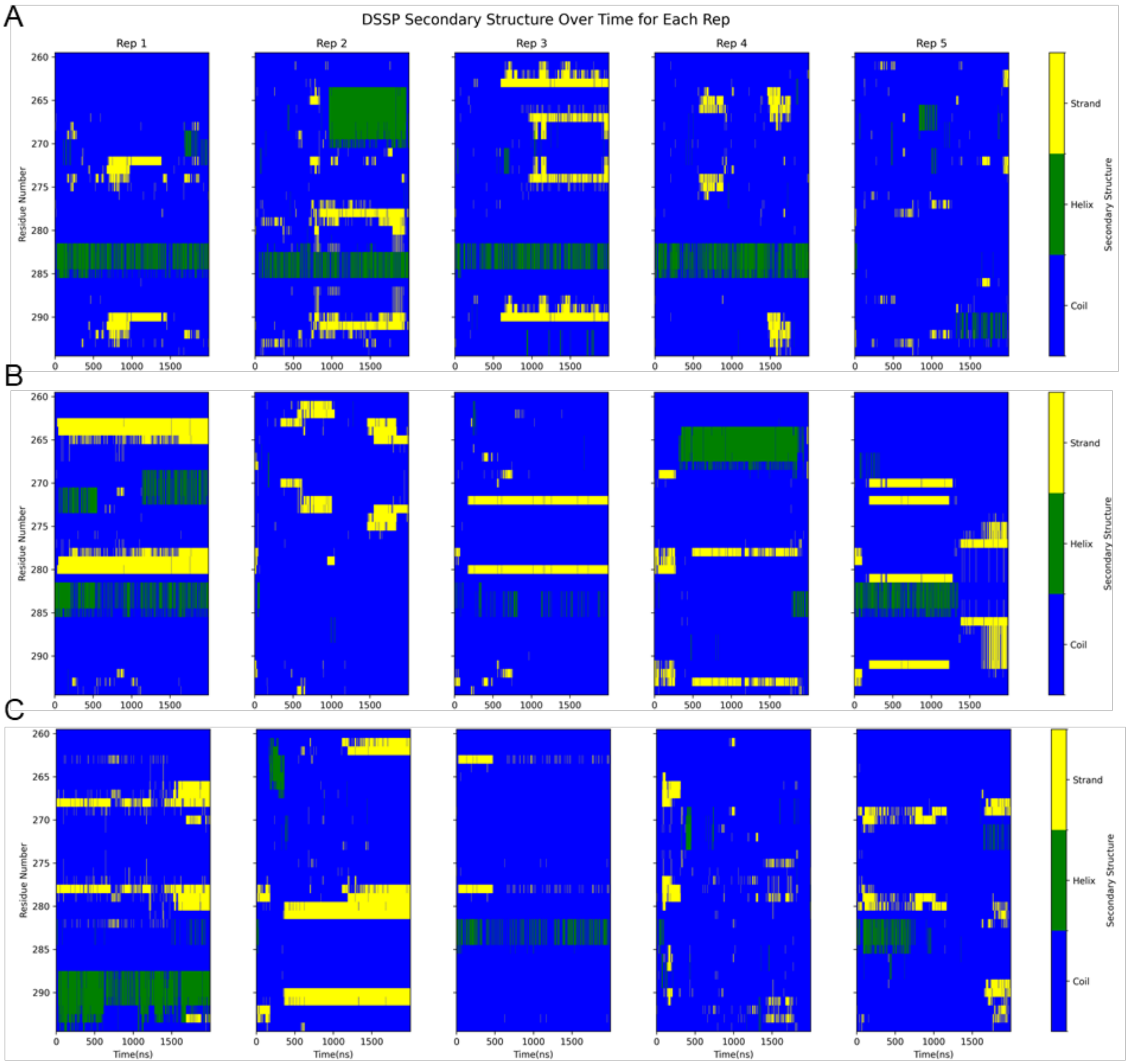


**Figure S16.** In MD simulations, the LASV FD can associate with membranes containing POPG and BMP, but not POPS. [A-C] Helical graph over the time course of the simulation for the different replicates indicates that the LASV FD has helical content in [A] POPC:POPG and [B] POPC:BMP, but not [C] POPC:POPS.

**Table S1.** Lipid composition of each membrane system utilized in molecular dynamics (MD) simulations.

| Membrane System | POPC (16:0, 18:1 (9Z)) | POPG (16:0, 18:1 (9Z)) | BMP (18:1, 18:1) | POPS (16:0, 18:1 (9Z)) | Total Lipid Count per Leaflet |
|---|---|---|---|---|---|
| POPC:POPG | 104 | 56 | 0 | 0 | 160 |
| POPC:BMP | 104 | 0 | 56 | 0 | 160 |
| POPC:POPS | 104 | 0 | 0 | 56 | 160 |

**Table S2.** The LASV FD is largely a random coil in the pre-fusion state. A three-state backbone code was assigned to each residue via TALOS+ to relate the NMR chemical shifts and backbone torsion angles: A – alpha, -160 < Φ < 0 and -70 < Ψ < 60; P – positive, 0 < Φ < 160 and -60 < Ψ < 95; B – beta, all others.

| Residue | A | P | B | Φ (deg) | Ψ (deg) | Classification |
|---|---|---|---|---|---|---|
| G260 | X | X | X | X | X | X |
| T261 | 0.3629 | 0.0445 | 0.5926 | 9999 | 9999 | B |
| F262 | 0.02 | 0.0038 | 0.9762 | -103.36 | 135.905 | B |
| T263 | 0.0072 | 0.0012 | 0.9916 | -124.138 | 135.929 | B |
| W264 | 0.0091 | 0.002 | 0.9889 | -140.552 | 160.013 | B |
| T265 | 0.0026 | 0 | 0.9974 | -90.04 | 142.794 | B |
| L266 | 0.1901 | 0.0053 | 0.8046 | -81.972 | 133.706 | B |
| S267 | 0.1238 | 0.0067 | 0.8695 | -87.453 | 131.924 | B |
| D268 | 0.9132 | 0 | 0.0868 | -62.05 | -37.185 | A |
| S269 | 0.9256 | 0.0081 | 0.0663 | -66.001 | -35.269 | A |
| E270 | 0.7835 | 0.0149 | 0.2016 | -99.847 | 2.97 | A |
| G271 | 0.0891 | 0.7638 | 0.1471 | 87.45 | 5.133 | P |
| K272 | 0.0459 | 0 | 0.9541 | -77.97 | 145.786 | B |
| D273 | 0.1483 | 0.0232 | 0.8285 | -90.186 | 133.528 | B |
| T274 | 0 | 0 | 1 | -87.326 | 126.207 | B |
| P275 | 0.1642 | 0.0092 | 0.8265 | -56.491 | 139.354 | B |
| G276 | 0.0379 | 0.9368 | 0.0253 | 71.27 | 13.882 | P |
| G277 | 0.0665 | 0.7898 | 0.1437 | 87.597 | 2.132 | P |
| Y278 | 0.0146 | 0 | 0.9854 | -72.939 | 140.26 | B |
| C279 | 0.5061 | 0.0253 | 0.4686 | -69.064 | -29.003 | A |
| L280 | 0.9162 | 0.0138 | 0.07 | -94.849 | -20.5 | A |
| T281 | 0.0559 | 0.0041 | 0.94 | -140.896 | 125.101 | B |
| R282 | 0.8469 | 0 | 0.1531 | -76.105 | -28.183 | A |
| W283 | 0.7979 | 0.0154 | 0.1867 | -72.422 | -30.108 | A |
| M284 | 0.8202 | 0.0143 | 0.1655 | -71.043 | -23.787 | A |
| L285 | 0.1161 | 0.0255 | 0.8584 | -75.178 | 130.323 | B |
| I286 | 0.0266 | 0 | 0.9734 | -97.401 | 137.845 | B |
| E287 | 0.1494 | 0.0082 | 0.8423 | -89.871 | 136.457 | B |
| A288 | 0.9317 | 0.0077 | 0.0606 | -70.208 | -26.748 | A |
| E289 | 0.2645 | 0.0476 | 0.688 | -103.589 | 147.751 | B |
| L290 | 0.0107 | 0.0056 | 0.9837 | -81.757 | 136.902 | B |
| K291 | 0.0281 | 0.0015 | 0.9704 | -100.366 | 146.75 | B |
| C292 | 0.0171 | 0 | 0.9829 | -87.151 | 116.576 | B |
| F293 | 0.3211 | 0.0255 | 0.6535 | -104.298 | 141.561 | B |
| G294 | 0 | 0.0031 | 0.9969 | -127.736 | 162.394 | B |
| N295 | 0.372 | 0.0498 | 0.5782 | 9999 | 9999 | B |

**Table S3.** Predicted dihedral angles suggest that the LASV FD has a helical structure within its FL in its post-fusion state. A three-state backbone code was assigned to each residue via TALOS+ to relate the NMR chemical shifts and backbone torsion angles: A – alpha, $-160 < \Phi < 0$ and $-70 < \Psi < 60$; P – positive, $0 < \Phi < 160$ and $-60 < \Psi < 95$; B – beta, all others. The residues likely to form a helix (black box) are located within the FL from $R^{282}$ to $L^{290}$.

| Residue | A | P | B | Φ (deg) | Ψ (deg) | Classification |
|---|---|---|---|---|---|---|
| G260 | 0.5068 | 0.0437 | 0.4495 | 9999 | 9999 | N |
| T261 | 0.0235 | 0 | 0.9765 | -96.604 | 138.83 | B |
| F262 | 0.3067 | 0.013 | 0.6804 | -101.298 | 129.364 | B |
| T263 | 0.1028 | 0.0218 | 0.8755 | -109.042 | 138.603 | B |
| W264 | 0.0032 | 0.004 | 0.9928 | -138.058 | 163.49 | B |
| T265 | 0.0019 | 0 | 0.9981 | -88.179 | 152.043 | B |
| L266 | 0.5564 | 0 | 0.4436 | -82.183 | 137.134 | A |
| S267 | 0.4401 | 0.0407 | 0.5192 | -66.043 | -34.703 | B |
| D268 | 0.9278 | 0.0036 | 0.0686 | -68.762 | -31.162 | A |
| S269 | 0.9466 | 0.0073 | 0.0461 | -65.386 | -34.876 | A |
| E270 | 0.9587 | 0 | 0.0413 | -97.485 | 0.763 | A |
| G271 | 0.0986 | 0.8361 | 0.0653 | 78.626 | 16.904 | P |
| K272 | 0.1452 | 0.012 | 0.8428 | -75.683 | 144.486 | B |
| D273 | 0.169 | 0.0318 | 0.7993 | -86.531 | 141.921 | B |
| T274 | 0 | 0 | 1 | -94.543 | 126.679 | B |
| P275 | 0.259 | 0.0147 | 0.7264 | -56.491 | 139.354 | B |
| G276 | 0.0389 | 0.943 | 0.0181 | 68.861 | 18.746 | P |
| G277 | 0.0694 | 0.8347 | 0.0959 | 86.458 | 1.243 | P |
| Y278 | 0.0216 | 0 | 0.9784 | -83.047 | 138.209 | B |
| C279 | 0.6291 | 0.0277 | 0.3432 | -80.975 | 115.565 | A |
| L280 | 0.0176 | 0.0008 | 0.9816 | -105.654 | 134.554 | B |
| T281 | 0.1842 | 0.0049 | 0.8109 | -86.045 | 142.567 | B |
| R282 | 0.9033 | 0 | 0.0967 | -69.85 | -31.059 | A |
| W283 | 0.9265 | 0.0195 | 0.0539 | -72.727 | -33.439 | A |
| M284 | 0.9522 | 0.0014 | 0.0464 | -66.117 | -36.429 | A |
| L285 | 0.987 | 0.0043 | 0.0088 | -78.996 | -30.413 | A |
| I286 | 0.8864 | 0.0063 | 0.1073 | -73.005 | -27.414 | A |
| E287 | 0.9854 | 0.0061 | 0.0085 | -67.92 | -37.857 | A |
| A288 | 0.9877 | 0.0033 | 0.009 | -64.826 | -33.075 | A |
| E289 | 0.9816 | 0.0047 | 0.0136 | -79.945 | -20.587 | A |
| L290 | 0.6746 | 0.016 | 0.3094 | -84.02 | -26.751 | A |
| K291 | 0.2883 | 0.0155 | 0.6962 | -101.144 | 146.926 | B |
| C292 | 0.2778 | 0.0155 | 0.7066 | -72.498 | -28.285 | B |
| F293 | 0.9382 | 0.0141 | 0.0476 | -100.604 | -7.119 | A |
| G294 | 0 | 0.0459 | 0.9541 | -121.627 | 146.372 | B |
| N295 | 0.3975 | 0.0624 | 0.5401 | 9999 | 9999 | N |

**Table S4.** Comparison of the average relative intensities of the LASV FD in different paramagnetic probes. In the pre-fusion state, both components of the FD have similar signals and are solvent-exposed. However, in the post-fusion state, the FP remains solvent-exposed, whereas the FL was inserted into the lipid head group of the membrane.

| | **PRE-FUSION** | **POST-FUSION** | | |
|---|---|---|---|---|
| | **Gd-DTPA** | **Gd-DTPA** | **5-DSA** | **16-DSA** |
| FD | 0.420 ± 0.021 | 0.222 ± 0.026 | 0.439 ± 0.037 | 0.620 ± 0.027 |
| FP | 0.458 ± 0.027 | 0.126 ± 0.022 | 0.530 ± 0.068 | 0.623 ± 0.046 |
| FL | 0.431 ± 0.025 | 0.317 ± 0.038 | 0.353 ± 0.034 | 0.620 ± 0.039 |

**Table S5.** Comparison of the overall relaxation times of the LASV FD in the different fusion states. In the pre-fusion state, the FP was slightly more flexible than the FL with values supportive of a random coil conformation. The FP continued to be more flexible than the FL in the post-fusion state, but to a much greater extent, with the FL becoming restricted.

| | **PRE-FUSION** | | | **POST-FUSION** | | |
|---|---|---|---|---|---|---|
| | **$^1H$ – $^{15}N$ NOEs (s)** | **$R_1$ ($s^{-1}$)** | **$R_2$ ($s^{-1}$)** | **$^1H$ – $^{15}N$ NOEs (s)** | **$R_1$ ($s^{-1}$)** | **$R_2$ ($s^{-1}$)** |
| FD | 0.448 ± 0.027 | 1.735 ± 0.091 | 4.941 ± 0.356 | 0.419 ± 0.038 | 1.305 ± 0.040 | 6.109 ± 0.683 |
| FP | 0.369 ± 0.018 | 1.750 ± 0.159 | 4.137 ± 0.550 | 0.253 ± 0.034 | 1.280 ± 0.036 | 4.426 ± 0.390 |
| FL | 0.497 ± 0.046 | 1.651 ± 0.062 | 5.708 ± 0.501 | 0.564 ± 0.048 | 1.339 ± 0.078 | 7.947 ± 1.224 |

**Table S6.** The percentage that a given conformer is populated is residue-dependent in both the pre- and post-fusion states. Notably, in the pre-fusion state, the signal for the b conformer of W$^{264}$ and a conformer of D$^{268}$ (bolded) overlap with each other and are not an accurate representation of the conformer populations. In turn, peak intensities in the backbone strips were used to justify the assignment of each population. Residues that do not occupy a given conformer are blacked out. The first (a), second (b), third (c), and fourth (d) most populated conformers of a given residue are shown.

| | | PRE-FUSION | | | | POST-FUSION | | | |
|---|---|---|---|---|---|---|---|---|---|
| | | a | b | c | d | a | b | c | d |
| FP | F262 | 53% | 37% | 10% | | 59% | 26% | 9% | 6% |
| FP | T263 | 76% | 13% | 11% | | 45% | 28% | 27% | |
| FP | **W264** | **34%** | **56%** | **10%** | | 70% | 13% | 9% | 8% |
| FP | T265 | 71% | 29% | | | 53% | 29% | 18% | |
| FP | L266 | 38% | 25% | 22% | 15% | 40% | 24% | 20% | 16% |
| FP | S267 | 79% | 21% | | | 68% | 32% | | |
| FP | **D268** | **78%** | **22%** | | | 100% | | | |
| FP | K272 | 91% | 9% | | | 88% | 12% | | |
| FP | T274 | 95% | 5% | | | 91% | 9% | | |
| | Y278 | 43% | 27% | 23% | 7% | 100% | | | |
| FL | C279 | 71% | 29% | | | 61% | 39% | | |
| FL | L285 | 77% | 23% | | | 100% | | | |
| FL | I286 | 47% | 35% | 18% | | 100% | | | |
| FL | E287 | 89% | 11% | | | 100% | | | |
| FL | A288 | 79% | 21% | | | 100% | | | |
| FL | E289 | 46% | 32% | 22% | | 83% | 17% | | |
| FL | L290 | 67% | 67% | 12% | | 100% | | | |
| FL | C292 | 100% | | | | 57% | 43% | | |
| | F293 | 100% | | | | 78% | 22% | | |
| | **Average** | **67 ± 19%** | **24 ± 12%** | **16 ± 6%** | **11 ± 6%** | **67 ± 17%** | **25 ± 11%** | **16 ± 8%** | **10 ± 6%** |

**Table S7.** In the pre-fusion state, the different conformers of the LASV FD have similar predicted dihedral angles. A three-state backbone code was assigned to each residue via TALOS+ to relate the NMR chemical shifts and backbone torsion angles: A – alpha, -160 < Φ < 0 and -70 < Ψ < 60; P – positive, 0 < Φ < 160 and -60 < Ψ < 95; B – beta, all others. The most populated conformer (a) is the residue label, whereas the second (b), third (c), and fourth (d) most populated conformers are indicated as such.

| Residue | A | P | B | Φ (deg) | Ψ (deg) | Classification |
|---|---|---|---|---|---|---|
| F262 | 0.0200 | 0.0038 | 0.9762 | -103.360 | 135.905 | B |
| F262b | 0.4000 | 0.0072 | 0.9528 | -99.874 | 146.588 | B |
| F262c | 0.0147 | 0.0016 | 0.9837 | -107.236 | 141.816 | B |
| T263 | 0.0072 | 0.0012 | 0.9916 | -124.138 | 135.929 | B |
| T263b | 0.0057 | 0.0015 | 0.9928 | -127.687 | 138.507 | B |
| T263c | 0.0233 | 0.0029 | 0.9738 | -108.678 | 131.182 | B |
| W264 | 0.0091 | 0.0020 | 0.9889 | -140.552 | 160.013 | B |
| W264b | 0.0096 | 0.0021 | 0.9883 | -140.353 | 158.993 | B |
| W264c | 0.0065 | 0.0023 | 0.9912 | -132.191 | 159.132 | B |
| T265 | 0.0026 | 0 | 0.9974 | -90.040 | 142.794 | B |
| T265b | 0.0029 | 0 | 0.9971 | -94.818 | 150.419 | B |
| L266 | 0.1901 | 0.0053 | 0.8046 | -81.972 | 133.706 | B |
| L266b | 0.2166 | 0.0018 | 0.7816 | -81.972 | 133.706 | B |
| L266c | 0.1909 | 0.0036 | 0.8055 | -81.972 | 133.706 | B |
| L266d | 0.1845 | 0.0027 | 0.8128 | -81.972 | 133.706 | B |
| S267 | 0.1238 | 0.0067 | 0.8695 | -87.453 | 131.924 | B |
| S267b | 0.2735 | 0.0140 | 0.7124 | -63.324 | -37.756 | B |
| D268 | 0.9132 | 0 | 0.0868 | -62.050 | -37.185 | A |
| D268b | 0.9444 | 0.0010 | 0.0546 | -64.607 | -35.129 | A |
| K272 | 0.0459 | 0 | 0.9541 | -77.970 | 145.786 | B |
| K272b | 0.0468 | 0 | 0.9532 | -79.519 | 136.657 | B |
| T274 | 0 | 0 | 1 | -87.326 | 126.207 | B |
| T274b | 0 | 0 | 1 | -115.991 | 127.654 | B |
| Y278 | 0.0146 | 0 | 0.9854 | -72.939 | 140.260 | B |
| Y278b | 0.0189 | 0 | 0.9811 | -72.939 | 140.260 | B |
| Y278c | 0.0034 | 0 | 0.9966 | -76.794 | 141.392 | B |
| Y278d | 0.0036 | 0 | 0.9966 | -77.226 | 143.301 | B |
| C279 | 0.5061 | 0.0253 | 0.4686 | -69.064 | -29.003 | A |
| C279b | 0.3114 | 0.0077 | 0.6808 | -101.077 | 142.122 | B |
| L285 | 0.1161 | 0.0255 | 0.8584 | -75.178 | 130.323 | B |
| L285b | 0.4407 | 0.0586 | 0.5007 | -69.516 | 126.270 | B |
| I286 | 0.0266 | 0 | 0.9734 | -97.401 | 137.845 | B |
| I286b | 0.0120 | 0.0056 | 0.9824 | -77.489 | 131.803 | B |
| I286c | 0.0445 | 0.0009 | 0.9546 | -96.844 | 150.896 | B |
| TABLE S7 CONTINUED ON NEXT PAGE | | | | | | |

| TABLE S7 CONTINUED | | | | | | |
|---|---|---|---|---|---|---|
| **Residue** | **A** | **P** | **B** | **Φ (deg)** | **Ψ (deg)** | **Classification** |
| E287 | 0.1494 | 0.0082 | 0.8423 | -89.871 | 136.457 | B |
| E287b | 0.0429 | 0 | 0.9571 | -90.715 | 144.977 | B |
| A288 | 0.9317 | 0.0077 | 0.0606 | -70.208 | -26.748 | A |
| A288b | 0.9257 | 0.0110 | 0.0633 | -75.261 | -21.553 | A |
| E289 | 0.2645 | 0.0476 | 0.688 | -103.589 | 147.751 | B |
| E289b | 0.1203 | 0.0278 | 0.8519 | -93.807 | 146.772 | B |
| E289c | 0.0893 | 0.0236 | 0.8871 | -87.517 | 134.364 | B |
| L290 | 0.0107 | 0.0056 | 0.9837 | -81.757 | 136.902 | B |
| L290b | 0.0155 | 0.0038 | 0.9808 | -80.729 | 134.134 | B |
| L290c | 0.0121 | 0.0045 | 0.9834 | -85.355 | 130.667 | B |

**Table S8.** Predicted dihedral angles of the different conformers are relatively similar for the LASV FD in the post-fusion state. A three-state backbone code was assigned to each residue via TALOS+ to relate the NMR chemical shifts and backbone torsion angles: A – alpha, -160 < Φ < 0 and -70 < Ψ < 60; P – positive, 0 < Φ < 160 and -60 < Ψ < 95; B – beta, all others. The most populated conformer (a) is the residue label, whereas the second (b), third (c), and fourth (d) most populated conformers are indicated as such.

| Residue | A | P | B | Φ (deg) | Ψ (deg) | Classification |
|---|---|---|---|---|---|---|
| F262 | 0.3067 | 0.0130 | 0.6804 | -101.298 | 129.364 | B |
| F262b | 0.2659 | 0.0236 | 0.7105 | -104.883 | 132.792 | B |
| F262c | 0.0752 | 0.0056 | 0.9191 | -102.444 | 131.969 | B |
| F262d | 0.0666 | 0.0052 | 0.9282 | -102.444 | 131.969 | B |
| T263 | 0.1028 | 0.0218 | 0.8755 | -109.042 | 138.603 | B |
| T263b | 0.0019 | 0.0049 | 0.9932 | -123.795 | 140.850 | B |
| T263c | 0.0291 | 0.0049 | 0.9660 | -111.131 | 141.192 | B |
| W264 | 0.0091 | 0.0020 | 0.9889 | -140.552 | 160.013 | B |
| W264b | 0.0095 | 0.0020 | 0.9885 | -139.437 | 159.124 | B |
| W264c | 0.0067 | 0.0027 | 0.9906 | -140.259 | 165.618 | B |
| W264d | 0.0069 | 0.0026 | 0.9905 | -140.259 | 165.618 | B |
| T265 | 0.0019 | 0 | 0.9981 | -88.179 | 152.043 | B |
| T265b | 0.0016 | 0 | 0.9984 | -92.295 | 154.232 | B |
| T265c | 0.0023 | 0 | 0.9977 | -94.363 | 154.151 | B |
| L266 | 0.5564 | 0 | 0.4436 | -82.183 | 137.134 | A |
| L266b | 0.3562 | 0 | 0.6438 | -82.183 | 137.134 | B |
| L266c | 0.8594 | 0 | 0.1406 | -68.489 | -24.282 | A |
| L266d | 0.8645 | 0 | 0.1355 | -68.489 | -24.282 | A |
| S267 | 0.4401 | 0.0407 | 0.5192 | -66.043 | -34.703 | B |
| S267b | 0.6084 | 0.0075 | 0.3841 | -64.255 | -33.783 | A |
| K272 | 0.1452 | 0.0120 | 0.8428 | -75.683 | 144.486 | B |
| K272b | 0.1128 | 0.0049 | 0.8823 | -74.603 | 134.747 | B |
| T274 | 0 | 0 | 1 | -94.543 | 126.679 | B |
| T274b | 0 | 0 | 1 | -119.516 | 126.546 | B |
| C279 | 0.6291 | 0.0277 | 0.3432 | -80.975 | 115.565 | A |
| C279b | 0.5678 | 0.0284 | 0.4038 | -75.107 | 122.862 | A |
| E289 | 0.9816 | 0.0047 | 0.0136 | -79.945 | -20.587 | A |
| E289b | 0.9869 | 0.0046 | 0.0085 | -79.557 | -19.575 | A |
| C292 | 0.2778 | 0.0155 | 0.7066 | -72.498 | -28.285 | B |
| C292b | 0.2577 | 0.0108 | 0.7315 | -72.498 | -28.285 | B |
| F293 | 0.9382 | 0.0141 | 0.0476 | -100.604 | -7.119 | A |
| F293b | 0.9430 | 0.0135 | 0.0436 | -104.646 | -3.126 | A |
